# Complex population history affects admixture analyses in nine-spined sticklebacks

**DOI:** 10.1101/2021.07.16.452636

**Authors:** Xueyun Feng, Juha Merilä, Ari Löytynoja

**Affiliations:** Organismal and Evolutionary Biology Research Programme, Faculty of Biological and Environmental Sciences, FI-00014 University of Helsinki, Finland; Area of Ecology and Biodiversity, Kadoorie Science Building, The University of Hong Kong, Pokfulam Road, Hong Kong, SAR; Institute of Biotechnology, HiLIFE, FI-00014 University of Helsinki, Finland

**Keywords:** Admixture, Introgression, Phylogeography, Sticklebacks

## Abstract

Introgressive hybridization is an important process in evolution but challenging to identify, undermining the efforts to understand its role and significance. On the other hand, many analytical methods assume direct descent from a single common ancestor, and admixture among populations can violate their assumptions and lead to seriously biased results. A detailed analysis of 888 whole genome sequences of nine-spined sticklebacks (*Pungitius pungitius*) revealed a complex pattern of population ancestry involving multiple waves of gene flow and introgression across northern Europe. The two recognized lineages were found to have drastically different histories and their secondary contact zone was wider than anticipated, displaying a smooth gradient of foreign ancestry with some curious deviations from the expected pattern. Interestingly, the freshwater isolates provided peeks into the past and helped to understand the intermediate states of evolutionary processes. Our analyses and findings paint a detailed picture of the complex colonization history of northern Europe and provide back-drop against which introgression and its role in evolution can be investigated. However, they also expose the challenges in analyses of admixed populations and demonstrate how hidden admixture and colonization history misleads the estimation of admixture proportions and population split times.

## INTRODUCTION

Population genomic analyses, such as inferences of selection or local adaptation, typically assume populations descending from a single shared ancestor. However, admixture of populations is common(Moran et al. 2021; Rius & Darling 2014), and multiple waves of migration and hybridization among divergent lineages can produce complex population histories (Flegontov et al. 2019; Hudson et al. 2021; Marques, Lucek, Sousa, Excoffier, & Seehausen 2019). Moreover, introgressive hybridization is recognized to contribute to adaptation and evolution-arynovelties(Hedrick 2013; Lamichhaney et al. 2018; Marques, Meier, & Seehausen 2019; Oziolor et al. 2019; Suarez-Gonzalez, Lexer, & Cronk 2018). The genetic signals of different evolutionary processes are hard to disentangle and, more seriously, unaccounted signals from complex admixture history may seriously mislead analytical methods designed for unadmixed data (Lawson, Van Dorp, & Falush 2018; Scerri et al. 2018). It is yet unclear how ancestral admixture events affect, for instance, the estimation of admixture proportions and divergence times among contemporary populations, potentially leading to misinterpretations regarding the reconstructed population histories (but see Flegontov et al. 2019).

Stickleback fishes are popular model systems for studying the genetic underpinnings of evolutionary changes in the wild. The well-developed genetic resources have made the three-spined stickleback(*Gasterosteus aculeatus*) a central model in evolutionary biology (reviewed in Reid, Bell, and Veeramah 2021), while the related nine-spined stickleback (*Pungitius pungitius*) has gained foothold as a model to study the genetic basis of local adaptations (e.g. Herczeg, Turtiainen, and Merilä 2010; Karhunen, Ovaskainen, Herczeg, and Merilä 2014; Kemppainen et al.2021; Wang et al. 2020). While both species occupy both freshwater and marine habitats, nine-spined sticklebacks commonly populate even the smallest freshwater ponds and lakes, making it an ideal model system for understanding the resilience of small populations under anthropogenic change. However, a detailed understanding of species’ history is central for comprehending the origins of adaptive and neutral variation, and the role of admixture and introgression as sources of genetic variation is widely acknowledged (Hellenthal et al. 2014; Hudson et al. 2021; Marques, Lucek, et al. 2019; Racimo, Sankararaman, Nielsen, & Huerta-Sánchez 2015).

Reconstruction of stickleback population histories has received a lot of recent attention (Aldenhoven, Miller, Corneli, & Shapiro 2010; Fang, Merilä, Matschiner, & Momigliano 2020; Guo et al. 2019; Mäkinen, Cano, & Merilä 2006; Marques, Lucek, et al. 2019; Orti, Bell, Reimchen, & Meyer 1994; Shikano, Shimada, Herczeg, & Merilä 2010; Wang et al. 2021). Earlier studies of nine-spined sticklebacks using microsatellites, mtDNA and single nucleotide polymorphisms (SNPs) derived with RAD-sequencing approaches (Bruneaux et al. 2013; Guo et al. 2019; Shikano, Shimada, et al. 2010; Teacher, Shikano, Karjalainen, & Merilä 2011) have revealed two distinct evolutionary lineages, the Western European lineage (WL) and the Eastern European lineage (EL), and their potential admixture in the southern parts of the Baltic Sea (BS) and around the Danish straits connecting the Baltic Sea basin to the North Sea and the Atlantic (Shikano, Shimada, et al. 2010; Teacher et al. 2011). However, the details of the colonization history especially in regard to admixture and its extent have not been worked out (Guo et al. 2019; Shikano, Shimada, et al. 2010; Teacher et al. 2011). Here, we tackled the question in unprecedented detail utilising a high-quality reference genome (Varadharajan et al. 2019), and calling SNPs for 888 whole-genome sequenced samples from 45 populations, covering a major part of the species’ circumpolar distribution range.

The primary aim of this study was to resolve the detailed phylogeographic history of a non-model organism by applying the latest population genetic methods to an extensive whole-genome sequence data. Given the likely secondary contact, we wanted to understand the extent and variability in admixture proportions among the populations across the contact zone. Since preliminary analyses revealed an unexpected population structuring, this motivated us to assess how the admixture and incorrect information about population history affects typical analyses of introgressive hybridization and population differentiation. We demonstrate that detailed assessments of population admixture are essential to our understanding of the evolutionary history of a lineage, and can help contextualise patterns of local adaptation, and even highlight the possible role of introgression in facilitating local adaptation.

## MATERIALS AND METHODS

### Sampling

The samples used in this study were collected in accordance with the national legislation of the countries concerned. A total of 888 nine-spined stickleback individuals (8–31 per population) were sampled from two previously identified European evolutionary lineages, EL (30 populations) and WL (10 populations), and five ancestral populations from Asia and North America. The species and lineage assignment of populations was based on information from previous studies (Guo et al.2019; Teacher et al. 2011) and was confirmed with data from this study (see Results). The samples were collected during the local breeding seasons with seine nets and minnow traps. After anaesthetising the fish with an overdose of MS-222, fin clips were preserved in 95% ethanol and stored at −80°C until DNA extraction. In addition, one *P. tymensis* individual serving as an outgroup was collected from Hokkaido, Japan (43°49’40”N, 145°5’10”E). The sampling sites are shown in Fig. 1; more detailed information, including sampling site coordinates and dates, sample sizes, population codes, and species names, is given in Table S1.

**FIGURE 1.**
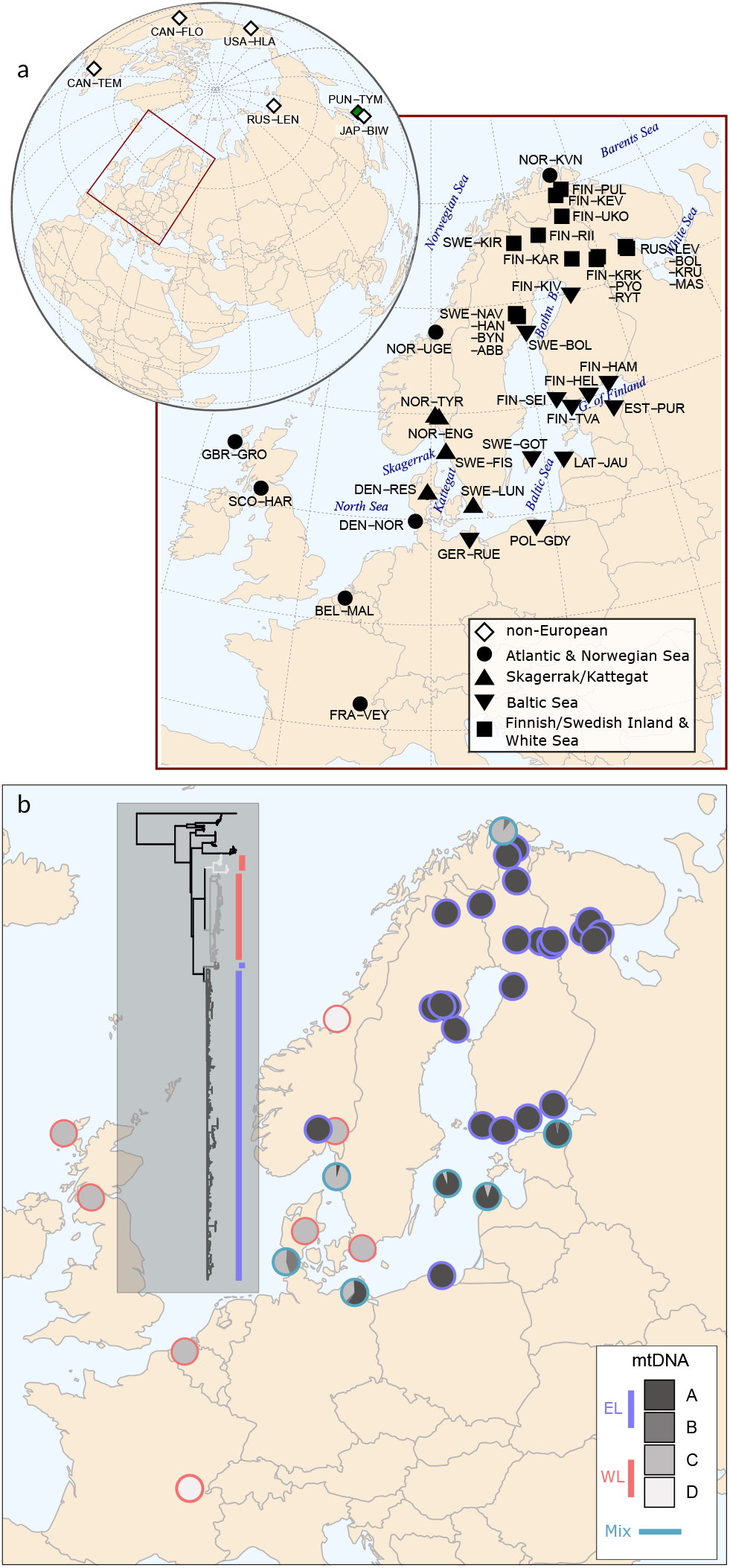
Sampling locations and European mtDNA lineages. a) 8-31 individuals per location were sampled, totalling 888 individuals. The shapes indicate the geographical origin of the sampled populations with filled shapes indicating the focal European populations, and white diamonds the reference populations from other parts of the world and green diamond the *P. tymensis* outgroup individual. b) The pie charts show the frequency of the four mtDNA types, indicated with the colors in the inset tree, for each population. The outline color (red, blue, cyan) indicates the mtDNA lineage assignment of the populations. The phylogenetic tree was constructed using RAxML on full mtDNA and rooted with *P. tymensis*.

### From sequencing to variant calling

Extractions of genomic DNA were conducted following the standard phenol-chloroform method (Sambrook & Russell 2006) from alcohol-preserved fin clips. DNA libraries with an insert size of 300–350 bp were constructed, and 150-bp paired-end reads were generated using an Illumina HiSeq 2500/4000 instrument. Library preparations and sequencing were carried out at the Beijing Genomics Institute (Hong Kong SAR, China) and the DNA Sequencing and Genomics Laboratory, University of Helsinki (Helsinki, Finland).

The reads were mapped to a subset of the nine-spined stickleback reference genome (Varadharajan et al. 2019) using the Burrows-Wheeler Aligner v.0.7.17 (BWA-MEM algorithm, Li 2013) and its default parameters. The subset, called V6b, contained 199 of the original 5,303 contigs and was 449 Mbp long; the removed contigs were inferred to be contamination, haplotypic copies or from the Y chromosome copy of LG12; a link to the included contigs can be found in the section Data and Code Availability. Duplicate reads were marked with SAMtools v.1.7 (Li et al. 2009) and variant calling was performed with the Genome Analysis Toolkit (GATK) v.4.0.1.2 (McKenna et al. 2010) following the GATK Best Practices workflows. In more detail, RealignerTargetCreator and IndelRealigner tools were applied to detect misalignments and realign reads around indels. The command HaplotypeCaller was used to call variants for each individual, parameters set as -stand_emit_conf 3, - stand_call_cof 10, -GQB (10,50), variant index type linear and variant index parameter 128000. The command GenotypeGVCFs was then used to call the variants jointly for all samples using default parameters. Interspecific variants were removed and binary SNPs were extracted with BCFtools v.1.7 (Li et al. 2009), excluding sites located within identified repetitive sequences (Varadharajan et al. 2019). Sites showing low (<8x) average coverage, low(<20) genotype quality and low(<30) quality score were filtered out using VCFtools v.0.1.5 (Danecek et al. 2011). For details of the subsequent filtering of the data sets used in different analyses, see Table S2.

### Analysis of population structure and phylogeny

Approximate population structure among all study samples was estimated using PCA within the PLINK toolset v.1.90 (Purcell et al.2007) and ancestry estimation within ADMIXTURE v.1.3.0 (Alexander, Novembre, & Lange 2009). In the latter, the analysis was replicated with the number of ancestral populations (K) varying from 2 to 4. In both PCA and ADMIXTURE analysis, variants with pairwise linkage disequilibrium above 0.1 (PLINK; –indep-pairwise 50 10 0.1) were removed.

The mtDNA variant calls were made haploid using the BCFtools plugin fixploidy. A maximum likelihood phylogeny was inferred using RAxMLv. 8.2.12 (Stamatakis 2014), under the GTRGAMMA model and using the lewis model to account for the ascertainment bias in SNP data (--asc-corr=lewis in RAxML). Branch support values were computed from 1,000 bootstrap replicates. The tree was rooted with *P. tymensis*.

For a phylogenetic analysis of nuclear DNA, two samples were randomly selected from each of the 45 *P. pungitius* populations and one *P. tymensis* sample was included as the root. The window size was set to 5,000 SNPs instead of physical length to have a similar signal for each local genomic region. Small windows at the chromosome ends and sites with more than 50% of missing data were discarded, leaving 1,792 valid regions for the analysis. Maximum likelihood phylogenies were inferred using RAxMLv. 8.2.12 (Stamatakis 2014) as for mitochondrial data, and the best tree for each region was selected from 20 alternative runs that started from different maximum parsimony trees (option -N 20 in RAxML). With the 1,792 best local trees, we then reconstructed a species tree under the multispecies coalescent model using ASTRAL v. 5.7.4 (Mirarab et al. 2014) with default settings. The Local Posterior Probabilities (Mirarab et al. 2014) were calculated for the main topology.

### Symmetry statistics and admixture tests

To get a global view of gene flow events, we started by computing the *f-branch* tests using Dsuite v.0.4r38 (Malinsky, Matschiner, & Svardal 2021) and the ASTRAL tree as the input topology. To elaborate the suggested gene flow events, the *f*- and *f4*-statistics were computed with ADMIXTOOLS v.5.1 (Patterson et al. 2012) using the programs qp3Pop and qpDstat with default parameters. The outgroup *f3*-statistics were computed in the form *f3*(pop1, pop2; outgroup) using qp3Pop. Outgroup-*f3* statistic measures the amount of shared drift between pop1 and pop2, higher *f3* values indicating closer affinity between pop1 and pop2 in comparison to the outgroup (see Fig. 4a), and allow determining the genetic similarity of populations. The f4 statistics (similar to D-statistics, Reich, Thangaraj, Patterson, Price, and Singh 2009) were computed in the form *f4*(Pop1, Pop2; Pop3, Pop4) using qpDstat. *F4* statistic measures the correlation of allele frequencies in the two pairs of populations and allows determining the existence of admixture. For f4 statistics, a deeply diverged Canadian population (CAN-TEM) was assigned as Pop4, while CAN-TEM and a Japanese marine population (JAP-BIW) were used as the outgroup in *f3* analyses; due to their greater resolution on European populations, *f3* results based on JAP-BIW were studied in detail. The heatmaps were generated with the R package gtools and the Neighbor-Joining tree from inverse distances using the R package APE.

**FIGURE 2.**
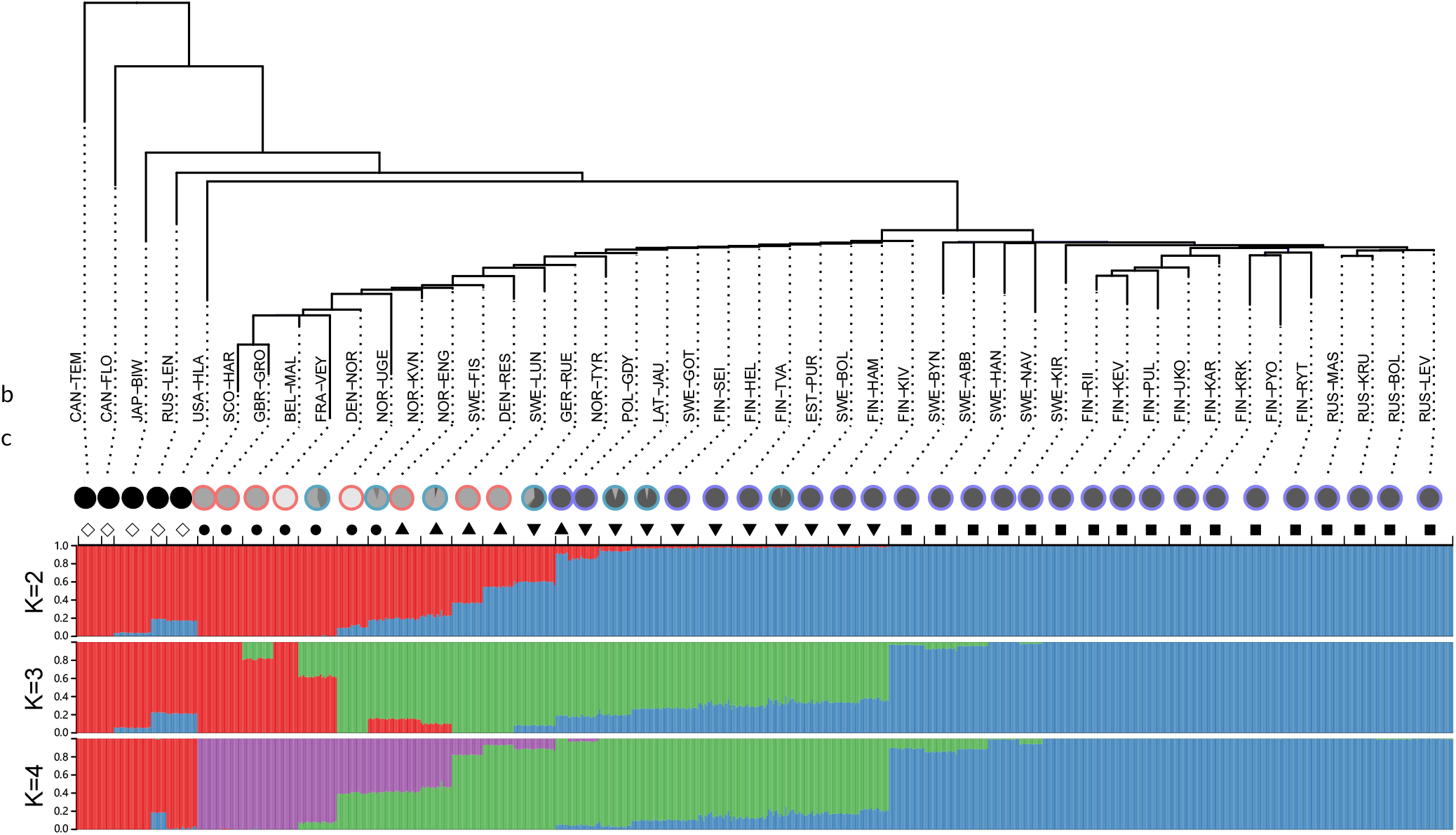
Genetic structure of European nine-spined stickleback populations. a) The nuclear phylogenetic tree was generated from 1,792 regional Maximum Likelihood trees using ASTRAL and rooted with *P. tymensis*. Two individuals were randomly selected from each population. For bootstrap values of each node see Fig. S2. b)The pie charts show the frequency of the four mtDNA types (see Fig. 1b)and the shapes indicate geographic origin of the sampled populations. Diamonds, filled circles, triangles, inverted triangles, and squares indicate non-European; Atlantic and Norwegian Sea; Skagerrak/Kattegat; Baltic Sea; and Finnish inland, Swedish inland and White Sea populations, respectively. c) The barplots show the proportion of genetic ancestry of the 888 whole-genome sequenced samples (x-axis) derived from K ancestral populations inferred with ADMIXTURE. K values >10 minimize cross-validation error but are uninformative about the deeper structure.

**FIGURE 3.**
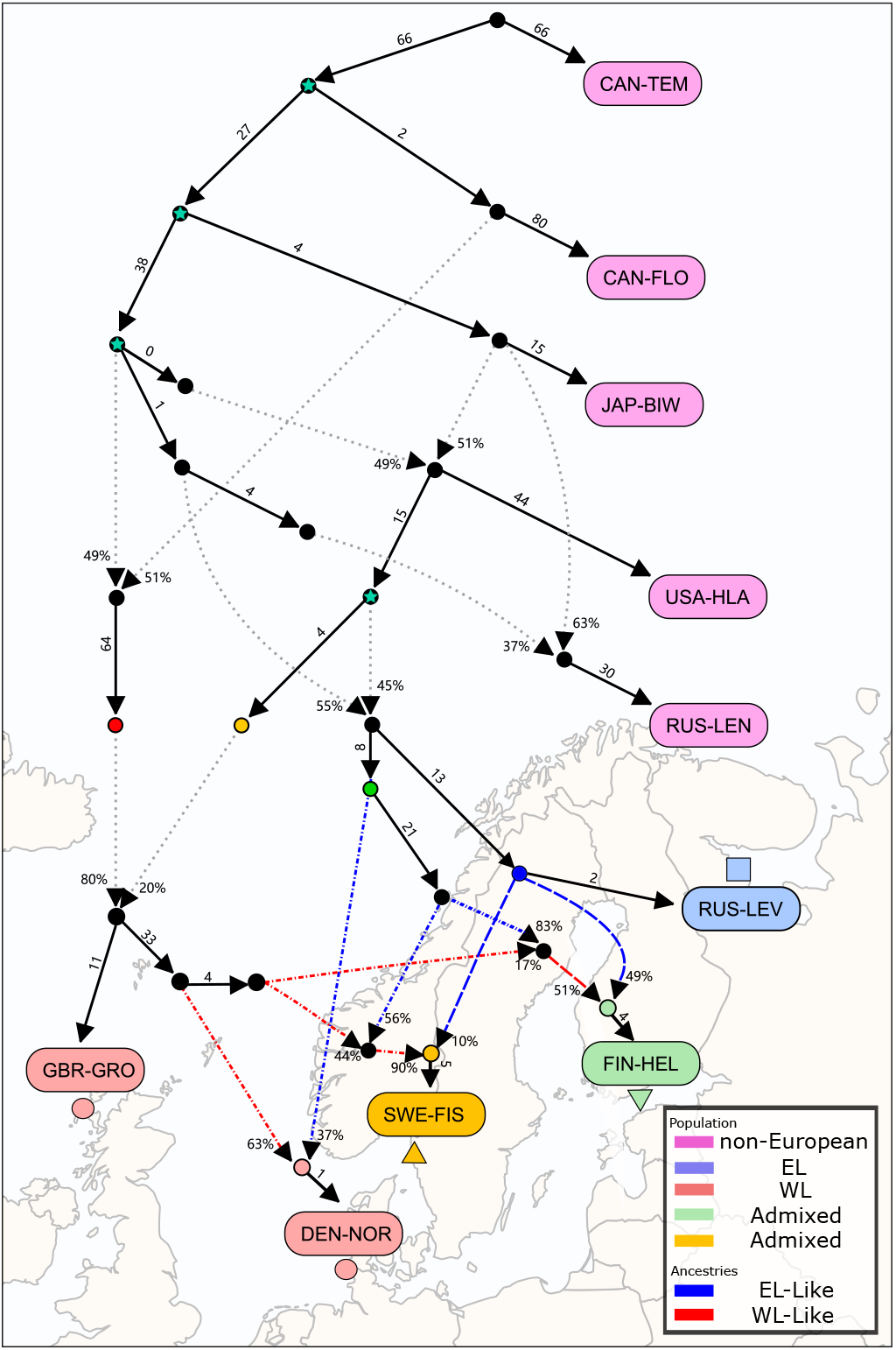
Complex history of European nine-spined stickleback lineages. The optimal qpGraph model (|Z|= 2.7) illustrates genetic drift and admixture events with solid and dotted/dashed lines, respectively, with the numbers indicating the amount (in units of *F_st_* × 1,000) and percentages. The UK population (GBR-GRO) descends 80% from a deep ancestor (red dot) but has a 20% contribution from an ancestral lineage (yellow dot) related to the White Sea population (RUS-LEV). The North Sea population (DEN-NOR) descends 63% from the old WL but also has a more recent 37% contribution from the east (green dot). The White Sea population (RUS-LEV) is a mixture of ancestral lineages forming the current day Japanese (JAP-BIW), Alaskan (USA-HLA) and the Lena river (RUS-LEN) populations. Populations from the Skagerrak(SWE-FIS) and the Baltic Sea (FIN-HEL) are mixtures of predominantly WL and EL origin, respectively. Red, yellow and green dots indicate hypothetical colonization waves to the Atlantic forming the current-day WL; blue dot indicates the ancestor of the EL that subsequently colonized the northern Fennoscandia. The stars in the graph (in cyan) mark the common ancestors of GBR-GRO and RUS-LEV.

**FIGURE 4.**
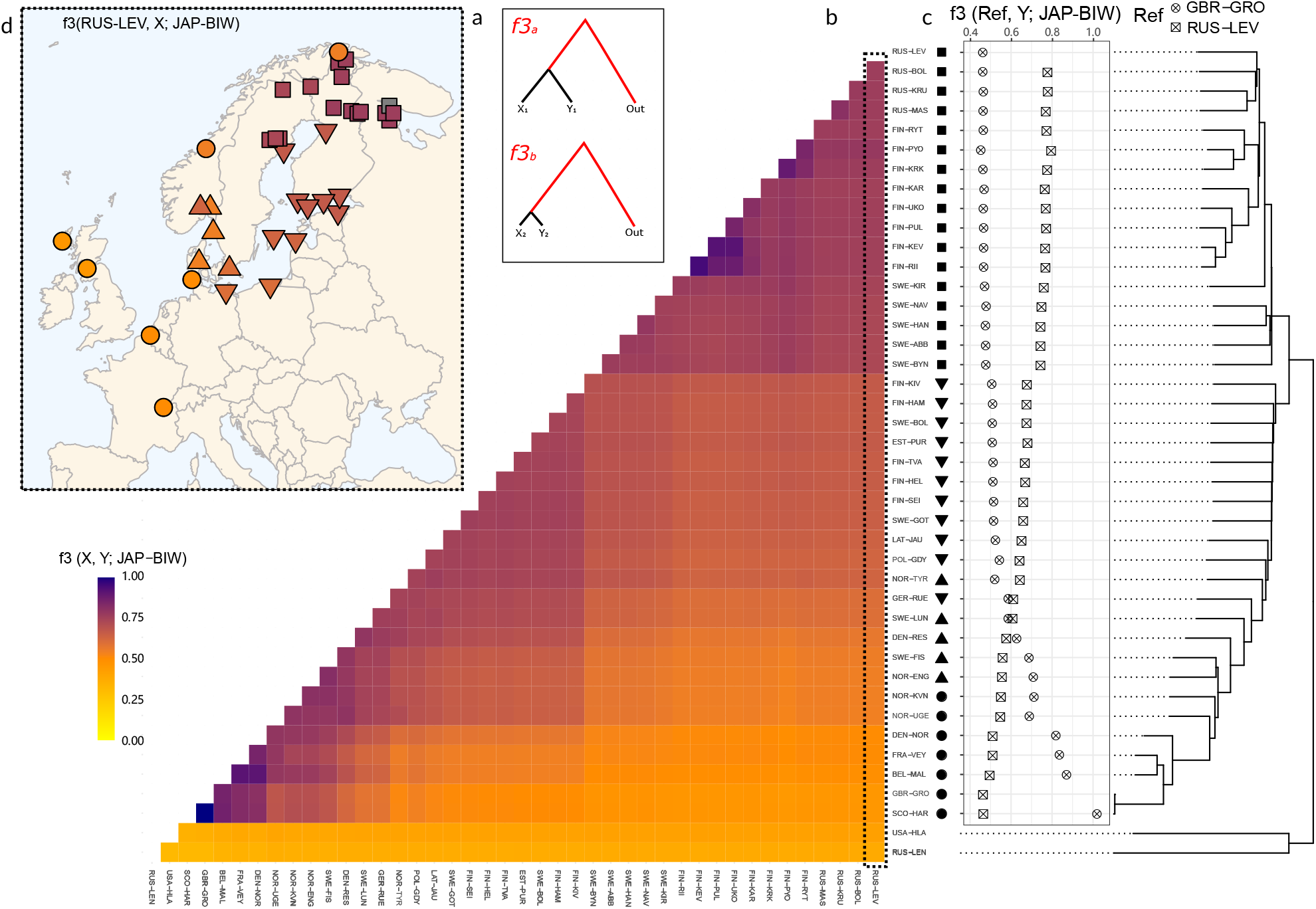
Relationships between WL and EL populations. a) A schematic example of outgroup-*f3* test with the branch lengths representing genetic drift. *X_1_* and *Y*_1_are distantly related and the relative length of the red outgroup branch is small (top); *X_2_* and *Y_2_* are closely related and the relative length of the red outgroup branch is large (bottom). The values *f3*(X, Y; Out) represent the length of the red branch for different pairs of *X* and *Y* with Out fixed as an outgroup. b) The heatmap colors show the statistics *f3*(X, Y; JAP-BIW), the darker values indicating closer affinity between populations X and Y. The shapes next to the population labels indicate different geographic areas and match those in the insert map. c) The symbols indicate the values *f3*(Reference, Y; JAP-BIW) for the White Sea (RUS-LEV; square with cross) and UK (GBR-GRO; circle with cross) populations. The NJ tree is based on the *f3*-statistics and is rooted with the non-European populations. d) The map shows the geographic location of the populations with shape colors indicating the values *f3*(RUS-LEV, Y; JAP-BIW) highlighted in the matrix. RUS-LEV is shown in grey.

The outgroup *f3*-statistics can be affected by genetic drift within the test populations (Patterson et al. 2012; Peter 2016), so we cross-validated our findings using the Rare-allele-sharing statistics (RASS; Flegontov et al. 2019). RASS shares similarity to f3-statistics but is driven by correlated frequency increases among rare alleles. Importantly, the SNP ascertainment is done using a fixed reference panel and thus is not affected by genetic drift among the test populations (Flegontov et al.2019). We defined the rare alleles using a panel consisting of the five ancestral non-European populations (RUS-LEN, USA-HLA, CAN-FLO, JAP-BIW, CAN-TEM; henceforth: ANC5) and representative populations from WL (UK; GBR-GRO) and EL (White Sea; RUS-LEV). Sites with more than 25% missing data were discarded, alleles were polarized using *P. tymensis* and, of the remaining sites, those with derived allele counts between 2-10 in the reference panel (in total 127 samples) were retained. For each frequency vector *x* and *y* (representing populations X and Y), sum of products (∑*_i_x_i_y_i_* is reported.

### Modelling admixture and estimation of admixture proportions

The minimum number of independent gene pools explaining the WL populations was inferred with the program qpWave (Reich et al. 2012) from ADMIXTOOLS v.5.1 (Patterson et al. 2012). qpWave relies on a matrix of f4 statistics, f4(left_1_,left_*i*_;right_1_, right_*x*_.), to infer the minimum number of streams of ancestries relating the “left” (target) populations to the “right” (source) populations. The p-values indicate whether the model with rank R is consistent with the data (p_rankR > 0.05); if the data are consistent, at least R+1 migration waves from the source populations are needed to explain the target populations (following Reich et al. 2012). We set the ANC5 populations and selected WL or EL populations as our “right” populations and inferred the minimum number of migration waves to different “left” populations (see Tables S3 & S6, SI Sections 1-2).

If a triplet (a group of three populations) can be modeled as derived from two sources of ancestry in qpWave, it follows that one of the populations can be modeled as a mixture of the other two populations with another tool, qpAdm (Flegontov et al. 2019; Harney, Patterson, Reich, & Wakeley 2021). To estimate the admixture proportions for the BS populations, we first used qpWave to find triplets (WL_*i*_, EL_*j*_, Baltic_*k*_) that show such patterns and then assessed the optimal source population pair based on qpAdm (Haak et al. 2015) analysis on other admixed populations (for details, see Fig. S8, SI Sections 1-2). The p-values given by qpAdm were used to assess the validity of each WL population, considering admixture models with p-value > 0.05 as plausible. Using the optimal WL and EL source populations, the relative proportions of WL/EL ancestry in the BS were quantified using f4-ratio (Patterson et al. 2012; Petr, Pääbo, Kelso, & Vernot 2019) and qpAdm (Lazaridis et al.2017). To elaborate the impact of unaccounted population structure on the estimation, WL populations with distinct population histories (i.e., with different numbers of migration waves) were used for estimation and comparison. The CAN-TEM and ANC5 populations were used as the outgroup for all f4-ratio and qpAdm analyses, respectively.

The complex relationships between diverse populations were modeled with qpGraph (Patterson et al. 2012). qpGraph assesses the fit of admixture graph models to allele frequency correlation patterns as measured by f-statistics. We started by fitting the deep ancestral populations and, using it as a basal model, gradually added different WL, EL and BS population combinations. Only graph models with the worst f4-statistic residual |Z|-score ≤ 3.5 were retained. For details of each qpGraph model, see SI Section 3.

### Sex chromosome (LG12) sequencing coverage

The ELand WL have different sex chromosome systems–heterogametic in EL (XY males) and dissimilar but yet unknown in WL (Natri, Merilä, & Shikano 2019)–and knowing the sex chromosome type may provide extra information about the origin of the admixed populations. In EL, chromosome LG12 consists of sex-chromosome and pseudo-autosome parts, allowing identification of EL males; in WL, the whole chromosome appears autosomal. We utilized the reference male (100X coverage) and five known females (each 10X) from the same FIN-PYO population (EL) to identify a set of sex-chromosome-associated SNPs. We first selected SNPs that had 5-15X coverage in each female and 30-75X combined coverage. We pruned down to 220,642 SNPs with coverage <50X in the male as the markers for the sex chromosome part in LG12 (region 1×10^7^-2×10^7^ bp), and to 430,056 SNPs with coverage >85X in the male as the markers for the pseudo-autosomal chromosome part in LG12 (region 2.75×10^7^-4.1×10^7^ bp). Finally, we computed the ratio of the mean ‘INFO/DP’ (approximate read depth across sample) across these regions and assigned ratios <0.6 as ‘Half’ and ratios >0.8 as ‘One’. Intermediate ratios and samples with less than 250,000 SNPs with coverage >5X in the pseudo-autosomal part were considered as ‘Unknown’. Due to atypical sequencing coverage, possibly due to the longer read length (251 nuc vs. 150 nuc in all other samples), the reference male showed the ratio of 0.77 and was assigned as Unknown. On the other hand, the other FIN-PYO samples showed expected ratios of 0.52-0.53 and 0.94-1.06 for males and females, respectively.

### Estimation of the divergence times

The population size histories for selected populations were inferred using MSMC2 v.2.1.1 (Malaspinas et al. 2016) with default parameters and assuming a mutation rate of 1.42×10^-8^ per generation (Guo, Chain, Bornberg-Bauer, Leder, & Merilä 2013) and generation length of 2 years (DeFaveri, Shikano, & Merilä 2014). Using these, we then inferred the divergence time for selected population pairs using the cross-population test implemented in MSMC2 (Malaspinas et al. 2016) following Schiffels and Wang (2020). The relative cross-population coalescent rate (CCR) value ranges between 0 and 1: CCR values close to 1 indicate that the two populations are one connected population, while the value of 0 means that the populations have fully separated. Following Schiffels and Durbin (2014), we consider a CCR of 0.5 as an indicator of population split. Demographic modelling with *moments*(Jouganous, Long, Ragsdale, & Gravel 2017) was used to infer the divergence time for the same population pairs. Following Sousa and Hey (2013) and Walsh et al. (2022), we tested five models: 1) strict isolation (SI) which assumes no migration between the two populations; 2) isolation with migration (IM) which assumes migration to be symmetric and constant; 3) two epochs (2EP) which allows one change in the migration rate; 4) secondary contact (SC) which assumes migration to have started after a period of isolation following the split of the two populations; and 5) ancestral migration (AM) which assumes migration at the early stages of divergence and strict isolation afterwards. Models with the highest average log-likelihood were chosen as the best model. Details of data processing, filtering and models for the two analyses can be found in SI Section 4, Table S2 and Script S1.

## RESULTS

The collected samples (Fig. 1a, Table S1) were whole-genome sequenced to 10-20X coverage, and variants were called using a standard pipeline. In total 15,217,577 SNPs across the 449 Mbp genome were included in the analyses after quality control (Table S2).

### Nuclear DNA reveals rampant admixture of ancestral lineages

The phylogenetic analysis of whole mtDNA data confirmed the earlier findings (Shikano, Shimada, et al. 2010; Teacher et al. 2011) and recovered a deep split between the WL and EL (Fig. 1b, Fig. S1). The Finnish and Swedish freshwater samples, the northern Baltic Sea samples and the White Sea samples all had the EL mtDNA type, whereas the Atlantic and Western European samples had the WL type (Fig. 1b). The EL type was found at low frequency in the Skagerrak, and the WL type on the German Baltic Sea coast and at low frequency among samples from Latvia and Gotland (Fig. 1b). In addition to clean EL and WL groups, the mtDNA phylogeny contained distinct early branchings at the root of both clades (Fig. 1b, Fig. S1). The mtDNA type branching from the WL clade was found with 100% frequency in isolated French (FRA-VEY) and central Norwegian freshwater (NOR-UGE) populations while the type branching from the EL clade was found in the North Sea (DEN-NOR), in two Baltic Sea populations (EST-PUR, GER-RUE) and at the Barents Sea coast (NOR-KVN; Fig. 1b). A clear discrepancy in the distribution of the main EL and WL types was Tyrifjorden (NOR-TYR), a lake outside of Oslo, Norway, that was of EL type with 100% frequency (Fig. 1b).

A nuclear phylogenetic tree was inferred from 1,792 regional Maximum Likelihood trees using ASTRAL (Fig. 2a, Fig. S2c; Mirarab et al.2014). Consistent with the mtDNA analysis, the nuclear tree grouped Fennoscandian and White Sea populations to EL (squares; Fig. 2) and Atlantic and Western European populations to WL (circles; Fig. 2). However, in contrast to mtDNA (Fig. S2), the nuclear data placed the Baltic Sea populations (inverted triangles; Fig. 2c) and several populations from the neighboring Skagerrak/Kattegat (triangles; Fig. 2c) basal to the WL clade with very short branch lengths (Fig. 2a, Fig. S2). Theoretical work on phylogenetic methods have shown that such patterns are expected for populations originating from admixture between distant lineages (Kopelman, Stone, Gascuel, & Rosenberg 2013; Rheindt & Edwards 2011), highlighting the challenges of using tree-based methods for analyses of population data. The possibility of a much wider admixture of the two major lineages was supported by population clustering with ADMIXTURE (Alexander et al. 2009) and Principal Component Analysis (PCA). In the ADMIXTURE plot, the intermediate populations showed a mixture of ancestral components associated with the EL and WL populations (Fig. 2c), and the PCA placed these populations as a continuum between the EL and WL populations (Fig. S3). Similarly to mtDNA analyses, the Tyrifjorden samples grouped among the BS populations (Fig. 2, Fig. S2).

### The Western lineage has complex history of admixture

A bifurcating tree cannot represent the relationships among admixed populations. To understand the extent of introgression, we performed an *f-branch* analysis with Dsuite (Malinsky et al. 2021) using the ASTRAL tree (Fig. 2a) as our phylogenetic model. While supporting the earlier findings of an EL clade and admixture in Skagerrak/Kattegat and BS populations, the *f-branch* analysis also unveiled a pattern of introgression from the ancestrally related Northeast Asian and North American populations to most WL populations and to Skagerrak/Kattegat/BS populations (Fig. S4). Notably, the absence of signal in the two UK populations suggests that the histories of WL populations are dissimilar and the non-UK populations have been affected by later pulses of migration.

To formally test the latter possibility, we used qpWave (Reich et al. 2012) and qpGraph (Patterson et al. 2012) to infer the history of migration waves. The qpWave analyses indicated at least three ancestry streams for the different WL populations (p_rank2 = 0.209, see SI Section 1, Tables S3-S6 and Reich et al. 2012). British, Belgian and French populations were inferred to have two streams of ancestry (p_rank1 = 0.228; Table S5) and the number increased to three when other WL populations (p_rank2 = 0.82 - 0.948; Table S5) were included (Table S5). The latter is of interest as the North Sea and central Norwegian freshwater population also carry multiple mtDNA types: the first of these required three streams of ancestry (p_rank2 = 0.972; Table S5) while with the latter the exclusion of the White Sea population (RUS-LEV) from the source set (“right” populations) led to decrease in the number of ancestry streams (p_rank1 = 0.0526; Table S5).

We applied qpGraph to build a model including the five early-branching Northeast Asian and North American populations (ANC5) as well as a lake population from the UK (GBR-GRO) and a White Sea marine population (RUS-LEV) as the representatives of the WL and EL, respectively. According to the best supported model (|Z|=1.6; Fig. S5), the UK population descends predominantly from two old waves from Asia but also has a minor contribution from a third wave shared with the EL. The White Sea population descends predominantly from an ancestor that itself was a mixture of lineages forming the current day Japanese and Alaskan populations (Fig. S5). Given this basal model, we added representative populations from the North Sea (Danish coast; DEN-NOR), the Skagerrak (SWE-FIS) and the Baltic Sea (Gulf of Finland; FIN-HEL) to the model. According to the best fitting model (|Z|= 2.7; Fig. 3), the Danish North Sea population is a mixture of the old WL (63%) and a later recolonization lineage (37%). The histories of the Skagerrak and Gulf of Finland samples are more complex: the Skagerrak has less contribution from the old WL than the North Sea population and has an additional 10% pulse of ancestry from the predecessor of the modern EL (blue dot; Fig. 3); in contrast, the Gulf of Finland is predominantly of EL origin and has approximately 9% contribution from the old WL forming the current day UK population (Fig. 3). Ancestry graphs are sensitive to changes in input data and often represent only one of many plausible arrangements (Lipson 2020; Patterson et al. 2012). While the choice of populations affected the order of branching, in our case the different and much deeper ancestry for the UK population was always strongly supported (Fig. S6).

### Secondary contact zone forms a gradient of foreign ancestry

To explore the population relationships in more detail, we computed the pairwise outgroup-*f3* statistics (Patterson et al. 2012). In this form, the *f3-statistic* measures the shared drift between two test populations in comparison to an outgroup population, the greater value indicating a closer affinity between the test populations (Fig. 4a). Using a Japanese population (JAP-BIW) as the outgroup, the *f3* analyses revealed major blocks reflecting the Atlantic WL populations, the pure EL populations, and a group of intermediate populations from the Baltic Sea and Skagerrak/Kattegat (Fig. 4b). The statistics also showed a finer grain gradient, with the more southern BS populations having closer affinity to WL and the more northern BS populations having closer affinity to EL (Fig. 4c,d). Among the WL populations, the two UK populations stood out and were different from all other populations (Fig. 4b). Repeating the analysis with our most distantly related outgroup population, CAN-TEM from Quebec, Canada, gave qualitatively similar but less resolved results (Fig. S7).

The qpWave and qpAdm analyses indicated that the North Sea and White Sea population are the optimal representatives of the WL and EL ancestral populations for studying the history of the admixed populations from the Baltic Sea area (see SI Sections 1-2, Fig. S8). We used the *f4*-ratio test (Patterson et al. 2012; Petr et al. 2019) to estimate separately the WL and EL ancestry proportions across the different populations (Fig. 5a,b, Tables S8-S9). Testing the WL ancestry, we saw it decreasing from 68% in the north-western extreme of the recognized contact zone (the Skagerrak) down to 22% in the south-eastern extreme of the area (the southern Baltic Sea); however, consistent with the earlier analyses, WL ancestry was seen in all BS populations, making 13-11% of the genome in the most distantly located populations (the Gulf of Finland and the Bothnian Bay; Fig. 5b). The EL ancestry mirrored this and decreased from 69% in the northern Baltic Sea to 45% in the southern parts, and further down to 25-20% in the Skagerrak (Fig. 5b). An alternative method, qpAdm (Lazaridis et al. 2017), gave consistent results (Fig. S8c).

**FIGURE 5.**
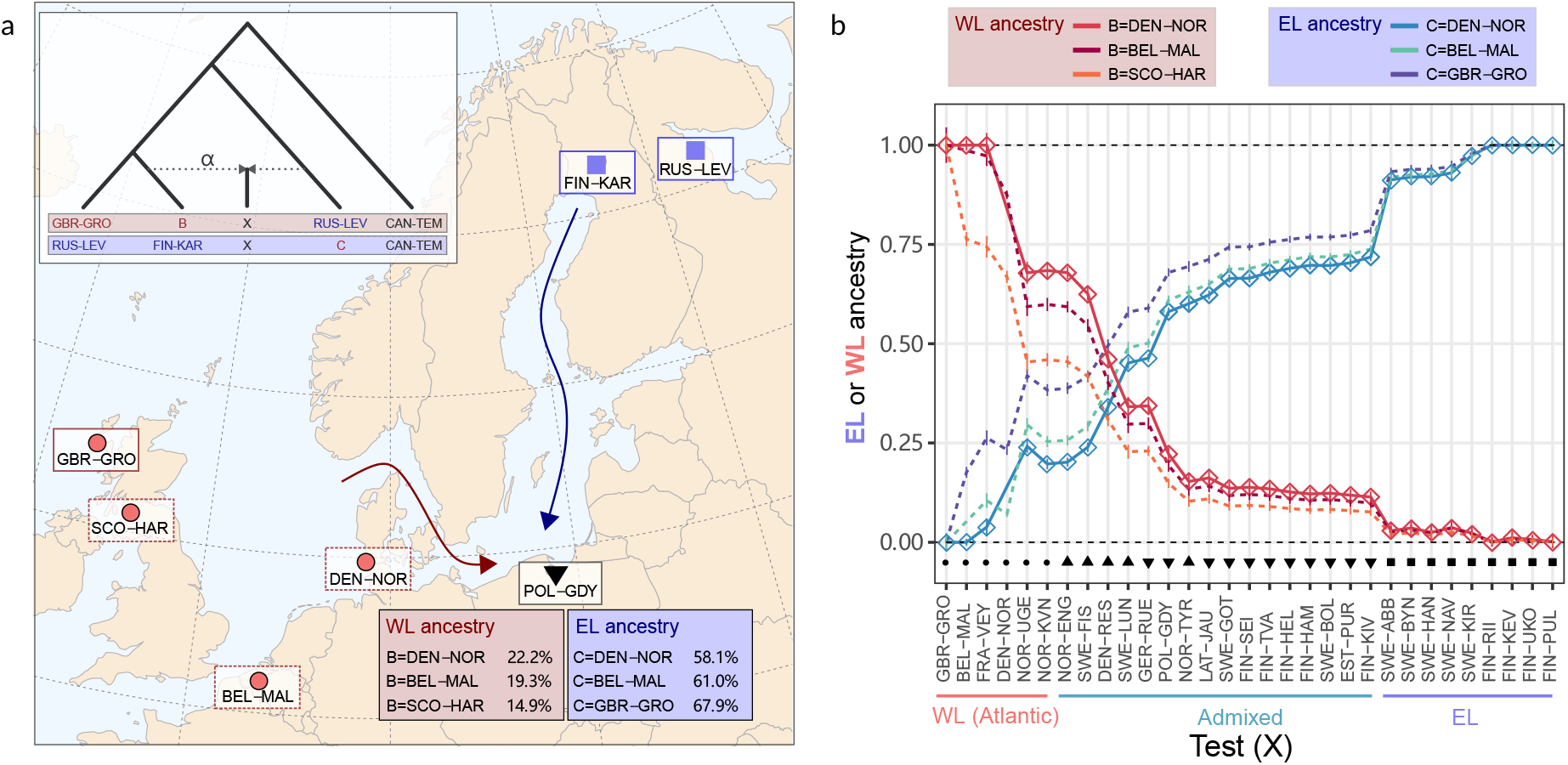
Quantification of WL and EL ancestry in admixed populations. a) The tree depicts the setup of the *f4*-ratio test for assessing the WL ancestry (red) and EL ancestry (blue). The test population X is assumed to be a mixture between populations B and C, *α* indicating the amount of WL (red) or EL (blue) ancestry. Here, the WL reference (GBR-GRO), the two EL populations (RUS-LEV, FIN-KAR) and the target population (X; PLO-GDY) are fixed, and the WL source (either B or C) change. This affects the outcome, and with different WL sources the admixed population is estimated to have 14.9-22.2% of WL ancestry and 67.9-58.1% of EL ancestry. b) Lines show the WL (blue shades) and EL (red shades) ancestry proportion (y axis) for different Test populations (x axis) using different WL source populations (colors). The estimates with DEN-NOR, the optimal WL reference amongst the studied populations, are indicated with diamonds. Vertical lines indicate 95% confidence intervals.

In the absence of the earlier analyses of ancestry and hisuorical adixture patterns, one could have acted with caution and selected a WL reference population further from the contact zone (e.g. UK populations). To test how the choice of reference affects the analyses, we recomputed the statistics using different populations as the WL reference (Fig. 5b; Tables S8-S10). Due to the complex structuring among the WL lineage, this indeed had a major impact and, with specific combinations of references, most WL populations showed significant f4-statistic and EL introgression (Fig. 5b, Table S10), naively interpreted to indicate recent gene flow out of the Baltic Sea. Using the UK populations as the WL reference, one would conclude that the North Sea has 23% of EL ancestry; while this could be geographically plausible, the analysis would also indicate that the recent EL gene flow has penetrated deep into eastern France, FRA-VEY showing 26% of EL ancestry. Recent EL migration to a small pond close to the Swiss border appears implausible and the result is more likely explained by an erroneous test setting and ancestral migration events being inferred as recent admixture (for more details, see SI 3).

To confirm that such a misleading signal can be created by an unaccounted ancestral admixture within WL, we simulated synthetic data using a population model that reflects the inferred history. While being a simplification of the real history (see SI Section 5 for more details), the simulations results nevertheless confirmed that the *f4*-ratio analysis is sensitive to unaccounted historical admixture among the populations (Patterson et al. 2012; Peter 2016) and the use of an incorrect reference population can inflate or deflate the estimates (Fig. S9).

### Admixture can seriously affect the divergence time estimation

The split time between the two ancestral lineages has been studied previously (Guo et al. 2019; Teacher et al. 2011) and, using calibration from geological events, was dated to be 0.67 Mya. To assess the impact of population structure on such estimates, we applied MSMC2 (Malaspinas et al. 2016) and demographic modelling with *moments* (Jouganous et al. 2017) on selected WL and EL populations. The cross-population analysis with MSMC2 showed a higher relative cross coalescence rate (CCR) between the White Sea and North Sea than between the White Sea - UK (Fig. 6a); consistent with this, the most recent split time was younger, 215,200 ya vs. 329,500 ya, respectively.

**FIGURE 6.**
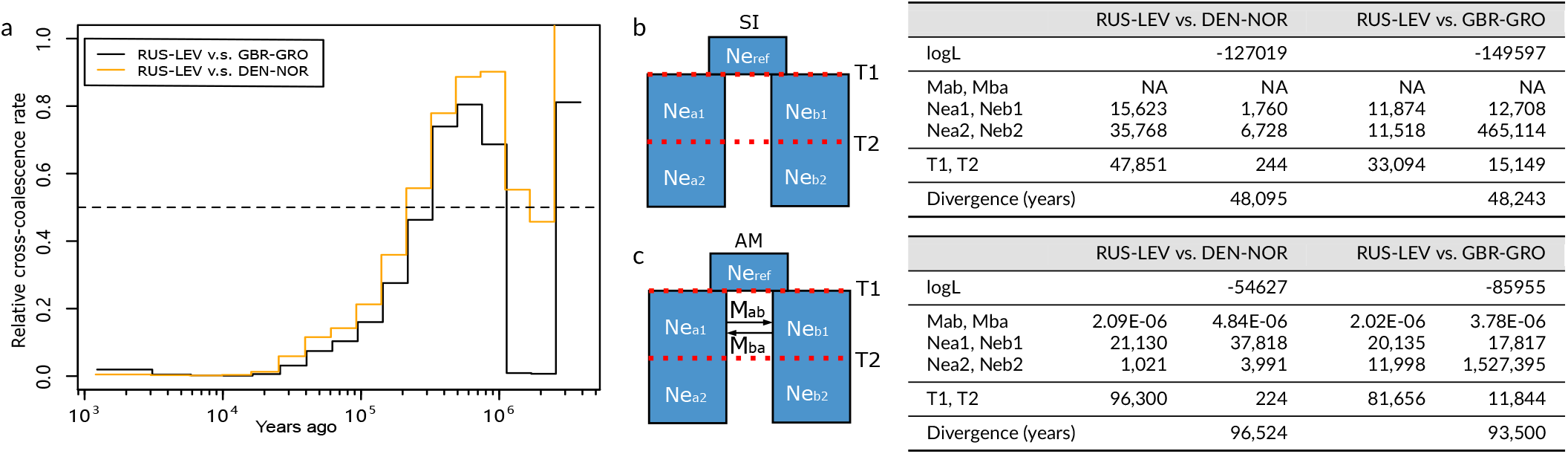
Dating of population separation times. a) Relative cross-coalescence rates for two population pairs estimated with MSMC2. Dotted lines indicate the time at which the relative cross-coalescence rate surpasses 0.5, considered here as the split time for the population pair. b) Schematic depiction of the strict isolation (SI) model used for the inference with *moments* (left), and the log likelihood and estimated parameters obtained (right). c) The ancestral migration (AM) model (left), and the log likelihood and estimated parameters obtained (right). Both models allow for one change in *N_e_*. Average values of the five replicates with the best likelihood are shown. Ne_ref, *N_e_* of ancestral population splitting into populations a and b; T1 and T2 (in years), time that *N_e_* of populations a and b remain constant as Nea1 & Neb1 for T1, and Nea2 & Neb2 for T2; T1+T2 is then the divergence time; Mab and Mba, migration rate from population a to b (Mab) and vice versa.

For demographic modelling, we built simple two-population models consisting of either presence or absence of migration and assuming constant population size (Fig. 6b). Unsurprisingly, the models with migration always fitted the data better and pushed the split times between the populations further back in time (Fig. 6c and Table S11). The estimated divergence time between the White Sea and North Sea/UK almost doubled when using the best model AM that allows for migration after the split event (Fig. 6c). Similar to our qpGraph model (Fig. 3), T2, the time period without gene flow, was higher between the White Sea and UK than between the White Sea and the North Sea, thus correctly capturing the differences in histories between the two WL populations. On the other hand, the divergence time estimates from *moments* were much younger than those from MSMC2, possibly reflecting the oversimplicity of the demographic models with limited population size changes.

We applied the divergence time estimation methods also for the synthetic data mimicking the inferred demographic history of the nine-spined sticklebacks (see SI Section 5). The analyses confirmed that the pulses of admixture push the cross-population coalescence ratio estimates further back in time, and that demographic models including migration give deeper divergence estimates (Fig. S12 and Table S12).

### Freshwater isolates provide windows to past admixture events

Peculiar aberrations in the ancestry analyses were the groupings of the four Norwegian freshwater populations (NOR-UGE, NOR-TYR, NOR-KVN and NOR-ENG) with geographically distant populations (Figs. 4 & 5). To confirm that the observed similarities are real and result from migrations after the latest secondary contact rather than from independent admixture events, we computed the rare allele sharing statistics (RASS; Flegontov et al. 2019). This method has been shown to be immune to even strong genetic drift (Flegontov et al. 2019) that often prevails in population isolates (Savolainen, Lascoux, & Merilä 2013). In addition, we exploited the information from the sex chromosomes, known to differ between the two evolutionary lineages (Natri et al.2019), to gain additional information of the population origins. However, the WL sex determination locus is unknown and we could only identify the EL males (the heterogametic sex), expected to show a sequencing coverage ratio close to 0.5:1 across the sex-chromosome and pseudo-autosomal parts of LG12 (Fig. S11, Kivikoski, Rastas, Löytynoja, and Merilä 2021).

In line with the *f4*-ratio and the outgroup-*f3* statistics, the two geographically closely located populations from Oslo were clearly differently related to EL and WL, with high levels of rare-allele sharing (RAS) seen between Tyrifjorden and BS populations (Fig. S10). Interestingly, samples from lake Engervann (NOR-ENG) - still connected to Skager-rak - showed higher RAS with central Norwegian isolated population than with the nearby Skagerrak samples (Fig. S10). The RAS patterns of samples from the Barents Sea coast and Skagerrak were nearly indistinguishable, strongly supporting that this Barents Sea coastal population is formed by recolonization from the North Sea/Skagerrak area rather than by an independent local admixture event (Fig. S10). Outside the contact zone, the presence of EL males matched the expected EL and WL ancestries of populations (data not shown). In the contact zone, exceptions to the expected pattern were the absence of EL males in the German Baltic Sea coast (GER-RUE) and the presence of them in central Denmark (DEN-RES), the southern tip of Sweden (SWE-LUN) and Tyrifjorden, Norway (Fig. 7). While seeing EL sex chromosomes in Tyrifjorden was not surprising, freshwater populations from central Denmark and Lund, Sweden showed a peculiar combination of EL sex chomosomes, WL mtDNA and predominantly WL nuclear DNA (Fig. 7). Although admixed nuclear DNA and mismatch between the sex chromosomes and mtDNA clearly indicate a mixed ancestry for the two populations, our sampling is too sparse to allow for an accurate reconstruction of the migration and admixture history.

**FIGURE 7.**
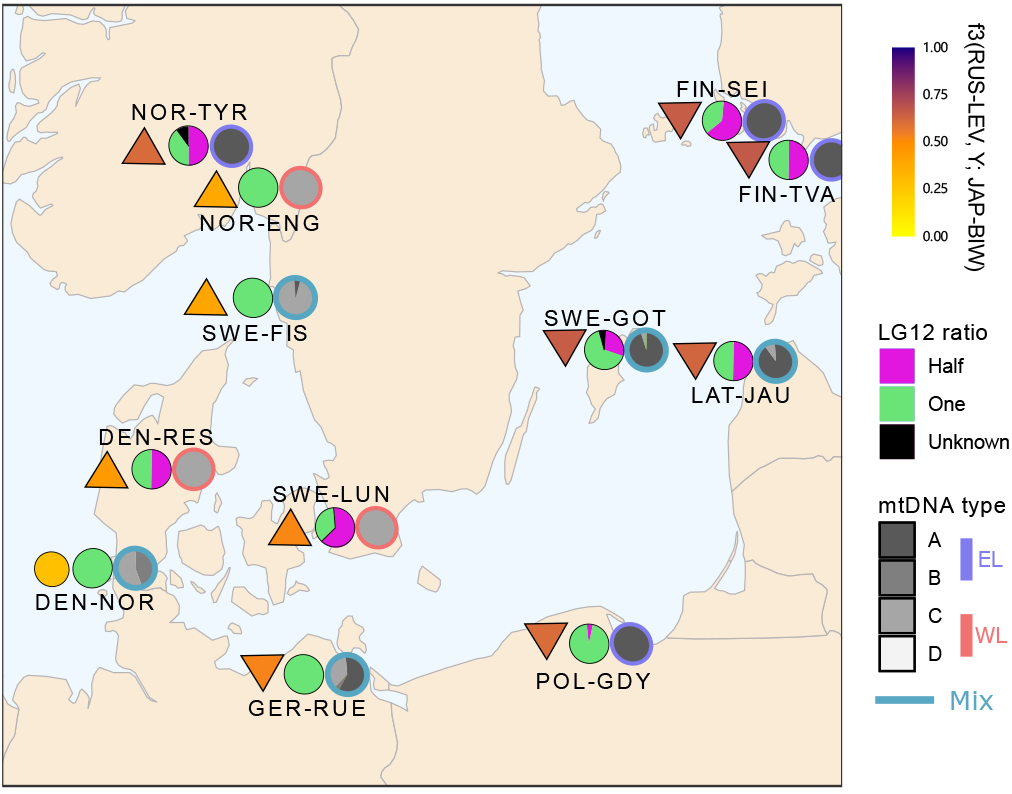
Conflicting genetic patterns in the secondary contact zone. For each population, the colored shapes show the outgroup-*f3* statistic (Fig. 4b), and the pie charts the LG12 and mtDNA (Fig. 2a) haplotype frequencies. LG12 ratio of “Half” indicates EL male whereas the ratio of “One” can be WL male or WL/EL female.

## DISCUSSION

An accurate picture of the historical relationships among contemporary populations is the foundation for all evolutionary inference. Due to climatic oscillations, multiple waves of colonization and admixture are common in the wild (Hudson et al. 2021; Marques, Lucek, et al. 2019; Schenekar, Lerceteau-Köhler, & Weiss 2014; Segawa et al.2021; Shirsekar et al. 2021) and many analytical methods can be seriously misled unless the resulting population structure and demographic history is correctly accounted for (Gompert & Buerkle 2016; Lange 2021; Scerri et al. 2018; Theunert & Slatkin 2017; Vitti, Grossman, & Sabeti 2013). The results of the current study provide a case in point: our analyses uncovered an unexpectedly complicated history of northern European sticklebacks pointing to multiple historical invasions and admixture events. They also showcase how admixture and introgression can influence present day patterns of genomic differentiation and our inferences on them. The results further illustrate how data from population isolates can be utilised to provide insights into the past evolutionary processes. In our case, the landlocked freshwater population isolates had preserved “genetic footprints” of historical events which helped us to piece together the complex history of the populations.

The presence of two diverged evolutionary lineages of European nine-spined sticklebacks is known from previous studies but their contact zone and admixture were thought to be limited to the Danish straits area (Guo et al. 2019; Teacher et al. 2011). Our results confirm the existence and admixture of the two lineages and show that the admixture zone is much wider than previously thought, spreading through the whole Baltic Sea and Skagerrak/Kattegat north of the Danish straits. Basic tools for population structure analyses, such as ADMIXTURE and PCA, were informative about the genetic ancestries of different populations, but failed to disentangle the complex history of migration contributing to the contemporary population structure. A reinterpretation of the results following outgroup-*f3*, *f-branch*, and qpWave analyses as well as modelling of the population history with qpGraph revealed a much more complex evolutionary history than originally anticipated. The history of the EL populations turned out to be rather simple, supporting recent radiation from an ancestral population in the White Sea area, with deeper relations to current day Japanese and Alaskan populations. In contrast, the WL populations were deeply structured and derived from multiple colonization events. In line with earlier hypotheses (Guo et al. 2019; Shikano, Shimada, et al. 2010; Teacher et al. 2011) of multiple waves of migration from Asia towards Europe, our analyses revealed that the WL and BS populations carry ancestries related to the Northeast Asian and North American populations. This component likely originates from an ancestral migration out of Asia to both Europe and North America (Guo et al. 2019) although a transAtlantic colonization has also been proposed (Aldenhoven et al. 2010). Whichever of these hypotheses is correct, our results support the view that at least three streams of ancestries, i.e. two old waves from Asia (or North America) and a third, more recent wave shared with the EL, have contributed to contemporary WL.

The results further showcase the importance of appropriate reference populations for reconstructions of population history and admixture proportions. Our estimates of ancestry proportions in the admixed populations changed drastically depending on the choice of reference populations. For instance, use of the UK population as the WL reference population seemed to suggest an “out of Baltic” event and the EL ancestry penetrating deeply into central France. However, replacing the UK population with the North Sea population, which was inferred to be closely related to the true WL ancestor for the admixed BS populations, this pattern disappeared. To confirm that our choice of reference populations is justified, we replicated the analyses with synthetic data reflecting the main events inferred to have taken place in the history of the WL populations. Although the simulation settings were a gross simplification of the real history, the results nevertheless supported the main findings: When the reference populations for the admixture analysis were more distantly related than the true parental populations, the ancestry from the other side (in our case from the EL) was overestimated. This highlights the challenges involved in tracking population history of admixed lineages: in order to avoid erroneous and biased inferences, comprehensive sampling and validation of potential source populations can be critical.

Our results also support the well known fact that admixture complicates estimation of phylogenies and divergence times (Hey & Nielsen 2004; Jones 2019; Long & Kubatko 2018; Pinho & Hey 2010). Classical phylogenetic trees are known to be unsuitable for the representation of histories of admixed populations (Chan et al. 2017; Harrison & Larson 2014; Supple, Papa, Hines, McMillan, & Counterman 2015). As a testimony of this, estimated nuclear phylogeny largely reflected the amount of shared ancestries amongst the populations and showed the typical symptom of admixed populations forming short branches intermediate to the two ancestral lineages (Kopelman et al. 2013; Rheindt & Edwards 2011).

To assess the impact of the population structure on inference of split times, we used two widely used approaches, the cross-population coalescence rate analysis with MSMC2 (Malaspinas et al. 2016) and the demographic modelling with *moments*(Jouganous et al. 2017). The results from empirical and synthetic data showed that both the gene flow and the choice of populations for the analysis as representative of the two evolutionary lineages had a major impact on the estimates. With *moments* analyses, the inclusion of migration always improved the fit of the data and doubled the inferred divergence times. Of the models tested, those with ancestral migration showed the best fit for modelling the history of WL and EL but the models behaved differently with the two WL populations studied, in line with the qpGraph model of multiple pulses of migrations into the WL. While these models are oversimplifications of the complicated history, they nevertheless illustrate the importance of accounting for changes in *N_e_* and historical migrations when dating the divergence time (Momigliano, Florin, & Merilä 2021). MSMC2 is poorly suited for the analysis of complex histories, but the estimated changes in cross-population coalescence rate correctly captured the extra pulse of gene flow inferred to have affected the North Sea population but not the UK population (Schiffels & Durbin 2014).

The divergence time estimation is highly sensitive to parameters used. We assumed a mutation rate of 1.42 × 10^-8^ per generation, originally estimated from transcriptome data with a divergence time of 13 Mya between the three- and nine-spined sticklebacks (Guo et al. 2013), and a generation length of 2 years (DeFaveri et al. 2014). Recent studies have pushed the divergence of the two species to 26 Mya (Varadharajan et al. 2019), thus potentially halving the mutation rate and doubling the estimated split times. A detailed determination of the WL-EL split is beyond the scope of this study and the simple analyses were primarily performed to assess the impact of gene flow on the estimates. In fact, even the definition of split time is unclear if the populations are connected by multiple waves of direct and indirect gene flow. Based on the results of the qpGraph model (Fig. 3), the most recent common ancestor for UK and White Sea was during the latest migration wave from the east and may have inhabited the Arctic Sea, while the earliest split between the lineages possibly happened hundreds of thousands of years earlier at the Pacific.

Our sampling of WL populations was sparse but we could still learn about the colonizations in different time scales. Isolation to ponds and lakes typically leads to increased drift and loss of genetic variation (Kemppainen et al. 2021; Savolainen et al. 2013; Shikano, Ramadevi, & Merilä 2010). Despite the smaller absolute amounts of variation, the isolates are of interest as they can provide insights into the early evolutionary history by preserving ancestral genetic variation. For instance, the central Danish freshwater population was found to have admixed nuclear DNA and Western mtDNA type but EL heterogametic sex chromosomes (this study and Natri et al. 2019) thus potentially representing the very early stage of the contact between two lineages around the Danish straits. In addition, our analyses indicate drastically different evolutionary histories for two lake populations only 15 km apart outside Oslo, Norway. The close affinity between Tyrifjorden and the Latvian river population (LAT-JAU) were confirmed with multiple alternative methods and we are confident that the signal indicates common ancestry, not independent admixture events resulting in similar ancestry proportions. A possible explanation for the Tyrifjorden population’s Baltic ancestry is the opening of the Baltic Sea basin at the end of the last ice age. In addition to connection through the Danish straits, there was a corridor across southern Sweden around 11,000 BP (Andrén et al. 2011). While this would have provided a relatively direct path from the central parts of the Baltic Sea towards North Sea, the timing may be too early for the ancestral Baltic Sea sticklebacks to be already admixed. On the other hand, the presence of gene flow from the Baltic Sea to Kattegat is supported by the Skagerrak marine populations showing low levels of EL ancestry.

It is also worth pointing out that many non-model systems lack source material (and resources) to study archaic DNA, limiting opportunities to study the ancestral steps of evolutionary processes. Access to the past states through population isolates as in the case of the nine-spined stickleback is a clear asset. Whatever the explanations for the aberrances in populations’ genetic and geographic identities are, they hint about the promise the isolated nine-spined stickleback lineages/populations carry for reconstruction of populations’ past history.

Reconstructing a more detailed history of colonization in the Baltic Sea area is beyond the scope of this study, but our results still provide some insights for further investigation. With the current estimates of mutation rate, the cross-population coalescence rate analysis of the Gulf of Finland and North Sea populations indicates that they have remained isolated for the last 25,700 years (Fig. S12b). If true, this means that there were nine-spined sticklebacks related to the current North Sea populations in the Baltic ice lake (12,600–10,300 BP; Leppäranta and Myrberg 2009), evolving independently from their Atlantic source, and thus the Baltic Sea basin was originally inhabited by WL populations and only later colonised, and largely taken over, by fish with EL ancestry. This is opposite to our initial thinking that the Baltic Sea is of EL origin and has only rather recently been introgressed by WL gene flow. A similar hypothesis of recolonization of the Baltic Sea from a southern refugium was proposed for the Atlantic salmon (Säisä et al. 2005). In line with this, we do find one extra pulse of ancestry in the southern Baltic Sea (Table S3). However, limited by our sampling, the origin of the Baltic Sea population is left unresolved and the timing of admixture may still be inaccurate. Although we could not directly estimate the ancestry contributed from the putative refugia population, our analyses suggest that its contribution is either small or largely represented by the North Sea ancestry (Shikano, Shimada, et al. 2010; Teacher et al. 2011). With data from relic isolates in the Baltics, Poland or Western Russia, a more detailed and complicated history could possibly be pieced together.

## CONCLUSIONS

In conclusion, the results indicate a very complex colonization and admixture history of northern Europe by nine-spined sticklebacks. While this is a case study of one particular species, it carries a broader relevance in a number of distinct ways. First, while earlier phylogeographic studies of these populations suggested fairly simple and straightforward scenarios to explain the current patterns genetic affinities among the different populations (Guo et al. 2019; Shikano, Shimada, et al. 2010; Teacher et al. 2011), our results suggests far more complex history involving repeated admixture events. Since many species have colonised the Atlantic Sea through the Artic Sea (e.g. Laakkonen, Strelkov, Lajus, and Väinölä 2015; Väinölä 2003; Vermeij 1991), it seems possible (if not likely) that similar complexities in evolutionary histories underline genetic relationships among populations of many other taxa too. Second, while multiple streams of ancestry in contemporary populations is acknowledged in increasing number of studies (Flegontov et al. 2019; Moreno-Mayar et al. 2018; Posth et al. 2018; Raghavan et al. 2015; Reich et al. 2012; Shirsekar et al. 2021; Skoglund et al. 2015), ancestral populations may also have been admixed or non-panmictic violating the assumptions underlying methods used to infer local adaptation, phylogenetic relationships among populations, as well as their divergence times. Hence, the most important lesson from this study is that when rampant admixture among multiple waves of colonization prevails, untangling the evolutionary histories of individual populations becomes extremely challenging and the underlying demographic models used to infer key population parameters (e.g. divergence time and admixture proportions) should be made with great caution (Momigliano et al. 2021).

## ACKNOWLEDGEMENTS

We thank Victor Berger, Pär Byström, Lasse Fast Jensen, Jacquelin De Faveri, Gabor Herczeg, Tuomas Leinonen, Scott McCairns, Andrew McColl, Heini Natri, Takahito Shikano and, Joost Raeymaekers for help in providing samples, and Miinastiina Issakainen, Sami Karja, Laura Häkkinen and Kirsi Kähkönen for help in the laboratory; Paolo Momigliano for help in *moments* anslyses. The advice and support from Antoine Fraimout, Baocheng Guo, Bohao Fang, Cui Wang, Paolo Momigliano, Pasi Rastas, Petri Kemppainen, Zitong Li, Mikko Kivikoski, Jarkko Salojärvi, Jack Beresford, Jacquelin De Faveri, Takahito Shikano, Simon Martin, and Martin Petr is gratefully acknowledged. Our research was supported by grants from the Academy Finland (# 129662, 134728 and 218343 to JM; # 322681 to AL), Helsinki Lifesciences Center (HiLife; to JM), China Scholarship Council (# 201608520032 to XF), and Finnish Cultural Foundation (#00210295 to XF). Computational resources provided by the CSC–IT Center for Science, Finland, are acknowledged with gratitude.

## AUTHOR CONTRIBUTIONS

J.M. started the project. X.F., J.M. and A.L. devised the research idea. X.F. performed the analyses with the help of A.L. X.F. and A.L. wrote the first draft and all authors participated in the writing of the final manuscript.

## DATA AND CODE AVAILABILITY

All the raw sequence data for this study can be accessed through European Nucleotide Archive (ENA) (https://www.ebi.ac.uk/ena) under accession code PRJEB39599. The list of contigs included in the reference genome version 6b as well as the code and scripts used in the analyses are available at https://github.com/XueyunF/nsp_phylogeo. Other relevant data (e.g. filtered VCF files, input and output files) are available from the Zenodo Open Repository: https://zenodo.org/record/6951309

