## Supplemental information for "Complex population history affects admixture analyses in nine-spined sticklebacks"

### Supplementary Information

1 Analysis of ancestry streams for WL, EL and BS populations . . . 2
2 Modelling of Baltic populations as a mixture of two lineages . . . 2
3 qpGraph modelling . . . 3
4 Estimation of divergence times with MSMC2 and moments . . . 3
5 Proof-of-concept data simulations and associated analyses . . . 3
6 Freshwater isolates provide windows to past admixture events . . . 4

### List of Figures

S1 Maximum likelihood tree on the mtDNA data . . . 5
S2 Cytonuclear incongruence . . . 6
S3 Principal Component Analysis of nuclear variant data . . . 7
S4 *f-branch* analysis of excess of shared derived alleles . . . 8
S5 qpGraph model for basal populations . . . 9
S6 qpGraph model for selected European populations . . . 10
S7 Outgroup-3 analysis with CAN-TEM as outgroup . . . 11
S8 Identification of optimal source pairs with qpWave and qpAdm . . . 12
S9 f4-ratio test with synthetic data . . . 13
S10 Rare Allele Sharing Statistics . . . 14
S11 Identification of EL males . . . 15
S12 Dating of population separation time with empirical and synthetic data . . . 16

### List of Tables

S1 Sample information, mtDNA type (A-D) and lineage assignment. . . . 17
S2 Data filtering for different analyses. . . . 18
S3 Detailed results of qpWave analysis: WL, EL and Baltic Sea populations. . . . 19
S4 Detailed results of qpWave analysis: WL and EL populations. . . . 20
S5 Detailed results of qpWave analysis: WL populations. . . . 21
S6 Two population symmetrical test with qpWave, the source was set as ANC5. . . . 22
S7 Detailed results qpWave testing of triples for the Baltic sea populations. . . . 22
S8 Detailed results of *f4*-ratio test with the EL as source. . . . 22
S9 Detailed results of *f4*-ratio test with the WL as source. . . . 22
S10 Detailed results of f4-statistics. . . . 22
S11 Detailed results of moments analysis for the empirical data. . . . 22
S12 Detailed results of moments analysis for the synthetic data. . . . 22

### List of Scripts

S1 Python-code for demographic analyses with *moments* . . . 23
S2 Python-code for data simulation with *msprime* . . . 23

#### SI 1 Analysis of ancestry streams for WL, EL and BS populations

We used qpWave (Reich et al. 2012) from ADMIXTOOLS (Patterson et al. 2012) to estimate the minimum number of streams of ancestry contributing to the WL, BS and EL populations. qpWave relies on a matrix of f4 statistics [f4(left$_1$,left$_i$;right$_1$, right$_x$)] to infer the minimum number of streams of ancestry relating the "left" (target) populations to the "right“ (source) populations. The p-values indicate whether the model with rank R is consistent with the data (p_rankR > 0.05); if it does, at least R+1 migration waves from the source populations are needed to explain the target populations (following Reich et al. 2012). The following set of populations were defined as "right" (source) populations for the analyses: CAN-TEM, CAN-FLO, JAP-BIW, RUS-LEN, USA-HLA; henceforth called ANC5.

Using ANC5 and RUS-LEV, a marine population from the White Sea, as the sources, we found the minimum number of migration pulses to be three for the nine WL populations (GBR-GRO, SCO-HAR, FRA-VEY, BEL-MAL, NOR-UGE, DEN-NOR, SWE-FIS, NOR-KVN and NOR-ENG; p_rank2 = 0.209, for details, see Table S3). According to the saturated model, two pulses originated from populations closely related to our Alaskan USA-HLA and central Canadian CAN-FLO, while the last pulse appeared to be from a population related to the current White Sea population RUS-LEV (Fig. 3). Importantly, the last pulse from RUS-LEV probably reflects a migration from a common ancestral population arriving from the east rather than introgression from the current EL populations. Using the same source set, we first tested the two British populations (GBR-GRO and SCO-HAR) and found the minimum number of migration pulses to be two (p_rank1 = 1, Table S4), with one of the likely sources being a population closely related to the Alaskan USA-HLA. When fixing the British populations in the target set and individually including the seven other WL populations, we found that only the populations from Belgium and central France did not show an increased rank (p_rank1 = 0.07-0.08, Table S4), indicating that they can be explained with the same two migration pulses. All the other WL populations showed significant change in the rank value (p_rank2 = 1, Table S4 and S5), meaning that these populations have received an additional pulse of migration which was not experienced by the other WL populations. The changes for the North Sea (DEN-NOR) and isolated mid Norwegian Sea (NOR-UGE) population (Table S5) are of interest as the two populations also carry early branching mtDNA types.

Using ANC5 and the British GBR-GRO as the source set, we quantified the number of ancestry streams for the 17 EL populations from the White Sea and the Finnish and Swedish ponds and lakes. We found the minimum number of streams to be four (p_rank3 = 0.828, Table S3). To elucidate the complex signature within EL populations, we performed two- and three-population qpWave tests. Of these, the White Sea (RUS-LEV/KRU/MAS/BOL) and the nearby Finnish pond populations (FIN-PYO/KRK/RYT/UKO/PUL) showed evidence of a single pulse (p_rank0 $>$ 0.05, Table S4) and symmetrical relationship to the outgroup set. Interestingly, the northern Finnish population FIN-KEV showed an additional pulse of gene flow (Table S4). The Swedish pond populations (except for SWE-HAN with three pulses) and the Finnish population FIN-KAR showed two pulses from the ancestral populations. This is consistent with these now isolated freshwater populations having at some stage been connected to the Baltic Sea basin and receiving WL-like ancestry from there.

Using ANC5, the British GBR-GRO and the White Sea RUS-LEV as the source set, we quantified the number of ancestry streams for the 12 Baltic Sea populations (FIN-KIV, FIN-HAM, SWE-BOL, EST-PUR, FIN-TVA, FIN-HEL, FIN-SEI, SWE-GOT, LAT-JAU, POL-GDY, GER-RUE, SWE-LUN). The minimum number of streams was found to be three (p_rank2= 0.071, Table S3). Removing southern Baltic sea populations (GER-RUE, POL-GDY, SWE-LUN, SWE-GOT) and the isolated Latvian population (LAT-JAU) decreased the migration waves to two (p_rank1=0.144, Table S3). This may indicate migration from a refugia population from the south into the Baltic sea (Gysels, Hellemans, Pampoulie, & Volckaert 2004; Teacher, Shikano, Karjalainen, & Merilä 2011).

#### SI 2 Modeling of Baltic populations as a mixture of two lineages

If a triplet can be modeled as derived from two sources of ancestry in qpWave, then one of the populations can be modeled as a mixture of the other two populations in qpAdm. Based on the phylogeny (Fig. 2b) and f4-statistic results (Table S10), the White Sea marine population RUS-LEV was chosen to represent the EL. We performed two-way admixture models in a group of three populations using qpWave (Reich et al. 2012) and qpAdm (Haak et al. 2015). First, using qpWave we identified triplets (WL$_i$, EL$_j$, Baltic$_k$) that can explain the Baltic Sea (BS) populations as derived from two ancestry streams (i.e. prank1_tail $>$ 0.05, Fig. S8a and Table S7). Second, the validity of the triplets and the chosen EL and WL populations as the optimal source pair was assessed based on qpAdm analyses of other admixed populations (Fig. S8b).

Given the results, the following populations were rejected as optimal WL source populations:

- The British Isle (GBR-GRO) and Scottish population (SCO-HAR) were rejected, as they strongly deviated from the rest of the WL populations and could not be used to distinguish the shared EL-like ancestry present in BS and other WL populations. Although they can be informative for studying the ancient genetic exchange between WL and EL, they are not a good proxy to study the recent admixture in the Baltic Sea.
- The central France (FRA-VEY) and Belgium (BEL-MAL) population were rejected, because 1) the two populations were found to be symmetrically related to GBR-GRO and SCO-HAR (Tables S4 & S6); and 2) several BS populations were rejected in the qpAdm analysis (Fig. S8b).
- The isolated Norwegian population (NOR-UGE) was rejected due to the rejection of SWE-FIS and NOR-ENG in the qpAdm analysis (Fig. S8b) despite strong evidence for their recent admixture.
- The freshwater populations DEN-RES and NOR-TYR have unique histories (see Discussion in the main text) and were expectedly rejected by qpWave and qpAdm.

Within our sampling, DEN-NOR was the best candidate for studying the WL ancestry in the BS populations:

- Using DEN-NOR as the WL source, the *f4*-ratio test (Fig. 5) correctly models FRA-VEY as un-admixed.
- The optimal qpGraph model (Fig. 3) indicates no recent EL gene flow to DEN-NOR.

Naturally, we cannot exclude the possibility of better proxies among unsampled populations.

#### SI 3 qpGraph modelling

We investigated the ancestry of different populations using modelling with qpGraph. In brief, for given allele frequency data and an input graph topology, qpGraph infers branch length and admixture proportions that minimize the differences between the observed and expected *f4*-statistics.

For the basal qpGraph model (Fig. S5), we used ANC5 as well as GBR-GRO (for WL) and RUS-LEV (for EL) to represent the two lineages. The fitted graph indicated that the Canadian lake population (CAN-FLO) is related to the British WL population. This can be a result of an ancient trans-Atlantic migration (Aldenhoven, Miller, Corneli, & Shapiro 2010), or derived from an ancestral population that colonized both North America and Europe (Guo et al. 2019). In line with the hypothesis of multiple waves of out-of-Asia migration, the eastern RUS-LEV and Alaskan USA-HLA populations were shown to derive over 50% of their ancestry from a lineage leading to Japanese JAP-BIW, and the rest from an ancestral population that is distantly related to the Japanese population.

Based on the basal model, we added one population from the North Sea (DEN-NOR), the Skagerrak/Kattegat (SWE-FIS) and the Baltic Sea (FIN-HEL). The three populations were chosen based on qpWave and qpAdm results to test if: 1) there was the North Sea – Baltic Sea connection; and 2) DEN-NOR is the optimal WL source population to model the admixed marine populations. Regardless of the order of populations being added, the model shown in Fig. 3 was the only optimal model.

Given the peculiar patterns observed in the NOR-UGE population, we set to test if it is indeed a relic population separated early in the admixture process. The optimal model found by adding NOR-UGE and DEN-NOR to the basal qpGraph model is shown in Fig. S6a. In this model, DEN-NOR and NOR-UGE share common ancestries but derive very different proportions of their genome from them: DEN-NOR has 62% of its ancestry from a WL-like ancestral population and only 38% from an EL-like ancestral population, while the numbers are nearly opposite, 33% and 67%, for NOR-UGE.

To explore the relationships among the WL populations, we excluded the ANC5 set and included the White Sea population (RUS-LEV) and the British Isles population (GBR-GRO) as representatives of the EL and WL, respectively, and added the four Atlantic WL populations into this model. The optimal model (Fig. S6b) shows that DEN-NOR and NOR-UGE received 53% and 31% of their ancestry from a common WL-like ancestral population. A descendant of this lineage then contributed to 61% of BEL-MAL and 55% of FRA-VEY ancestry. On the other hand, NOR-UGE and FRA-VEY inherit 69% and 45% of their ancestry from a EL-like population; this evolved further and contributed 47% and 39% of ancestry in DEN-NOR and BEL-MAL. The model is congruent with other lines of evidence showing that NOR-UGE and FRA-VEY are of earlier origin than the other (here, DEN-NOR and BEL-MAL) WL populations.

#### SI 4 Estimation of divergence times with MSMC2 and moments

We used MSMC2 to infer the split time between populations GBR-GRO, DEN-NOR, RUS-LEV and FIN-HEL. Sample specific VCF and mask files were first created of aligned reads (in bam format) using bamCaller.py from msmc-tools (available at https://github.com/stschiff/msmc-tools) and applying the mappability mask of Kivikoski, Rastas, Löytynoja, and Merilä 2021. The variants were then phased with SHAPEIT v.4.2 (Delaneau, Zagury, Robinson, Marchini, & Dermitzakis 2019) and the input for MSMC2 was finalised using generate_multihetsep.py from msmc-tools, following the instructions in Schiffels and Wang (2020). After the generation of the input data, two individuals from each population were randomly selected for analysis with MSMC2. Each individual was first analyzed separately and then used in the cross-population analysis to infer the split time between the paired population. The results were visualized with R scripts (from msmc-tools) assuming a mutation rate of 1.42e-8 and a generation length of 2 years.

We used *moments* (Jouganous, Long, Ragsdale, & Gravel 2017) to infer the divergence time under five models: 1) strict isolation (SI) which assumes no migration between the two populations; 2) isolation with migration (IM) which assumes migration to be symmetric and constant; 3) two epochs (2EP) which allows one change in the migration rate; 4) secondary contact (SC) which assumes migration to have started after a period of isolation following the split of the two populations; and 5) ancestral migration (AM) which assumes migration at the early stages of divergence and strict isolation afterwards. We first used ANGSD v.0.921-3-g40ac3d6 (Korneliussen, Albrechtsen, & Nielsen 2014) to estimate the folded site frequency spectrum (SFS) and then generated two-dimensional site frequency spectrums (2dSFS) for the selected population pairs. For each of the population pairs, the models were optimized in 10 independent runs, each consisting of 6 rounds of optimization and 30 replications for each optimization. We selected the best 10 replicates from each of the models according to the log-likelihood and estimated the divergence times. The same mutation rate and generation length were used as in MSMC2 analyses. For details, see Table S2 and Script S1.

#### SI 5 Proof-of-concept data simulations and associated analyses

We assessed the effect of complex population histories on the estimation of admixture proportions and divergence times using simulations. We generated synthetic data with *msprime* v.0.7.4 (Kelleher, Etheridge, & McVean 2016) using a population model that reflects the main events inferred to have taken place in the history of European nine-spined sticklebacks (see Script S2 and Fig. S9a). More precisely, we simulated an early WL colonization followed by a contact with the second-wave populations, mimicking the UK populations in our study (#1 / "GBR-GRO"); and one of the second-wave populations having a long independent history (#2 / "FRA-VEY"), while the other had an additional contact with the first WL population (#3 / "DEN-NOR"). The latter population then provided a pulse of admixture to a bottle-necked EL population, mimicking our Baltic Sea populations (#4 / e.g. "FIN-HEL"), while two other EL populations remained clean (#5, #6 / e.g. "FIN-KAR", "RUS-LEV"). We simulated 25 replicates, each 10 Mbp long, and sampled 20 diploids per population. The simulated data were used to replicate the analyses with *f4*-ratio, msmc2 and *moments*.

For MSMC2 and *f4*-ratio tests, we followed the same pipeline as for empirical data. For *moments* analyses, we first used vcf2dadi.py script (available at https://github.com/CoBiG2/RAD_Tools/blob/master/vcf2dadi.py) to generate the 2dSFS for each of the population pair and then analysed these with the same pipeline as the empirical data.

While the simulation settings were a simplification of the real history, the results nevertheless revealed that the *f4*-ratio analysis is sensitive to unaccounted historical admixture among the populations (Patterson et al. 2012; Peter 2016) and the use of an incorrect reference population can inflate or deflate the estimates. The use of the correct reference population, #3 ("DEN-NOR") in our analysis (Fig. S9b,c), gave 1-$\alpha$ estimates very close to the true simulated value of 25% (Fig. S9c). When using the more distantly related population #1 ("GBR-GRO") as the "WL" source in the *f4*-ratio test, we saw also populations #2 ("FRA-VEY") and #3 to obtain high 1-$\alpha$ values (Fig. S9c) despite no recent gene flow from "EL" was simulated (the gene flow from #3 to #4 was unidirectional; Fig. S9a).

In the MSMC2 analyses, the impacts of gene flow are captured by the most extreme population pairs: The split of populations #4 and #5, mimicking the admixed Baltic Sea population and its parental EL population, is accurately represented by the cross-population coalescence time jumping from 0.1 to 0.75 around 5000 generations ago (Fig. S12); however, the line does not reach 1, reflecting the deep variation obtained through admixture from the WL. On the other hand, when using 0.5 cross-population rate as the criterion, the deepest simulated divergence between populations #1 and #5, mimicking the earliest WL population and the parental EL population, is underestimated. Moreover, the line deviates from 0 at 6,000 years ago, correctly reflecting the more recent gene flow between the populations (Fig. S12).

Demographic modeling with *moments* demonstrated the impact of gene flow (Fig. S12 and Table S12): Strict Isolation (SI) gave much younger split times than models with migration. Modelling migration correctly improved the model fit (log-likelihood) and the estimated divergence time (Table S12). For populations with extra pulse of migration, the 2EP model always outperformed the IM model and gave the correct divergence estimates. On the other hand, unrealistic parameter estimates were seen when gene flow scenarios were incorrectly modeled.

### SI 6 Freshwater isolates provide windows to past events

Our sampling of WL populations was sparse but we could still learn about the colonizations in different time scales. As the WL has received multiple waves of gene flow from the east, we expected NOR-KVN, a population from the coastal Barents Sea in northern Norway, to show an intermediate position between the more southern WL populations and the EL. However, NOR-KVN showed close affinity to marine populations from the North Sea and Skagerrak and likely represents a recolonization from the south. On the other hand, NOR-UGE, an isolated lake population from western Norway, was found to have a slightly higher proportion of EL ancestry than the marine neighbors NOR-KVN and NOR-ENG (Fig. 5). This seems geographically implausible, and we believe that NOR-UGE is a representative of an early state of the latest mixing between the WL and EL lineages; due to its isolation, it has not been affected by the later admixture/migration events and thus provides untainted (but possibly fuzzy) signal not distinguishable in the more connected populations. By definition, the Rare Allele Sharing statistic (RASS) focuses on rare alleles and NOR-UGE shares those with all other WL populations. While lake isolates such as NOR-UGE may be strongly affected by drift and selection, adding multiple independent relic populations to the analyses in future should provide additional cues about the complex evolutionary history of these fish.

One of the peculiar findings of our analyses was the drastically different evolutionary histories of two populations outside Oslo, Norway. Engervannet (NOR-ENG) is a small pond connected by a 0.5 km long stream to Oslofjord and Skagerrak, and the nine-spined stickleback population there was genetically similar to the nearby marine populations. On the contrary, Tyrifjorden (NOR-TYR), an inland lake some 15 km west of Engervannet, had a nine-spined stickleback population most similar to a freshwater population from Latvia (LAT-JAU; see the main text). While the history of NOR-TYR is enigmatic, other nearby freshwater populations may have even more complex histories. DEN-RES, a population from Silkeborg, Denmark, shares many rare alleles with the nearby SWE-FIS and SWE-LUN populations, but even more so with NOR-UGE, an isolated pond population from midwest Norway. In RASS, the top five populations for DEN-RES also include LAT-JAU from the Baltic Sea and NOR-KVN from northern Norway. In addition to admixed nuclear DNA, DEN-RES was found to share western mtDNA but EL sex chromosomes. The fact that the two ancestral lineages have different sex chromosome systems (Natri, Merilä, & Shikano 2019) makes the admixture process and the history of contact zone isolates especially intriguing. However, it should be noted that EL and WL have been shown to produce fertile offspring (Natri et al. 2019), and the divergent sex determination system does not disqualify any of our findings.

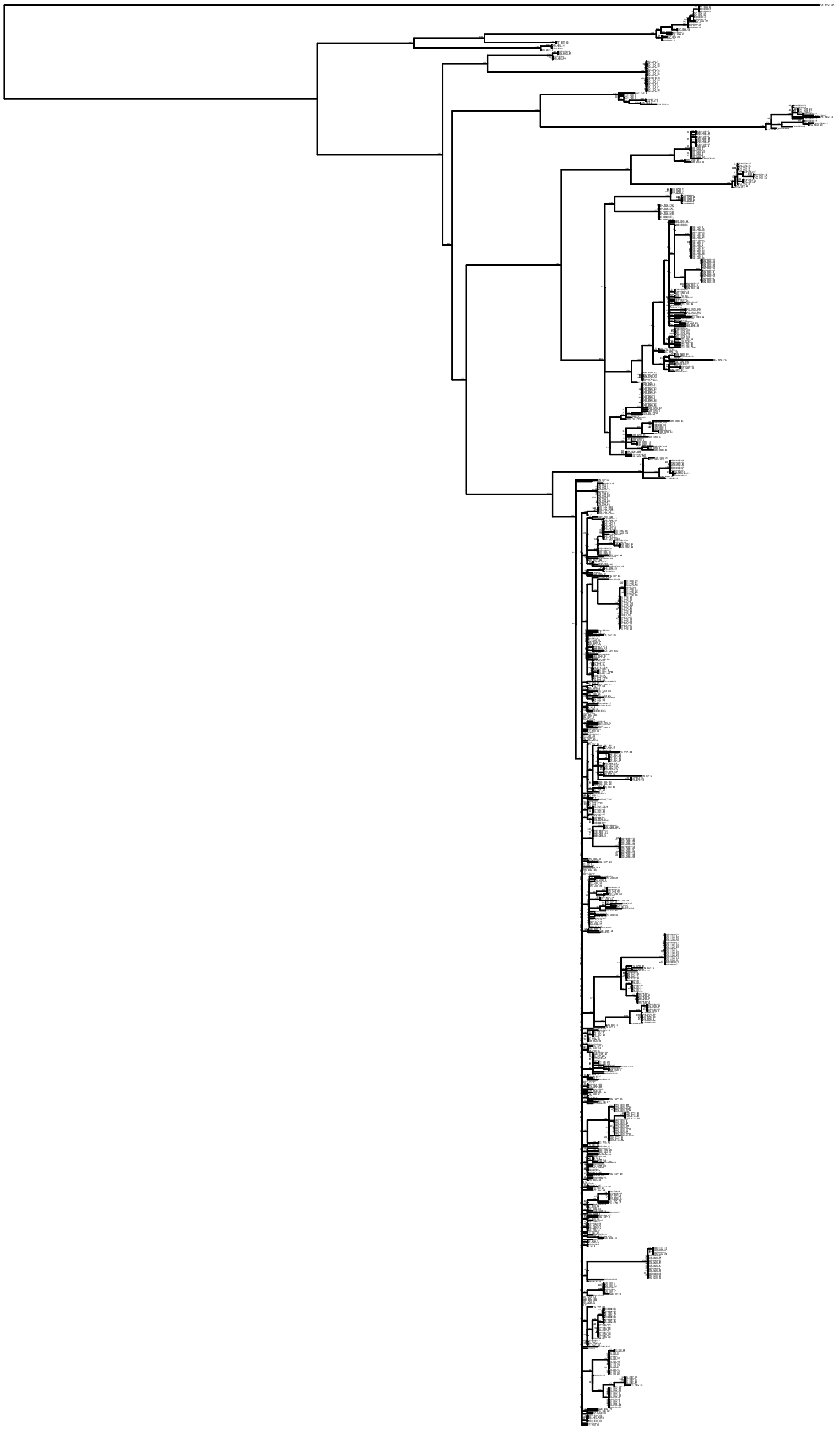

**FIGURE S1** The RAxML maximum likelihood tree for the mtDNA data with the taxon names and bootstrap support values.

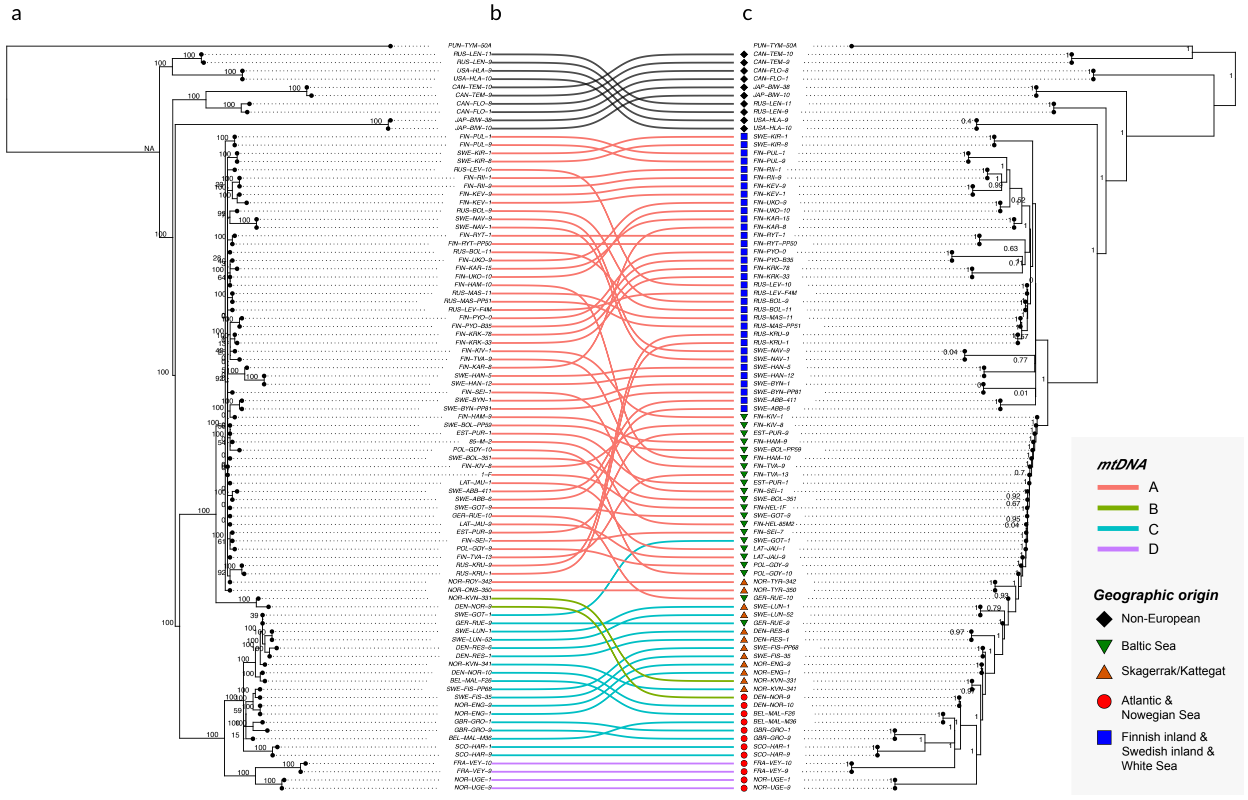


**FIGURE S2** Cytonuclear incongruence. a) The RAxML maximum likelihood tree for mtDNA data with taxon names and bootstrap support values. b) Lines, colored by the mitochondria type (cf. Fig. 1b), show incongruences between the two phylogenetic trees. c) The ASTRAL tree for autosomal DNA data with taxon names and local posterior probabilities. The colored shapes indicate the geographical origin of each sample and match those in Fig. 2b.

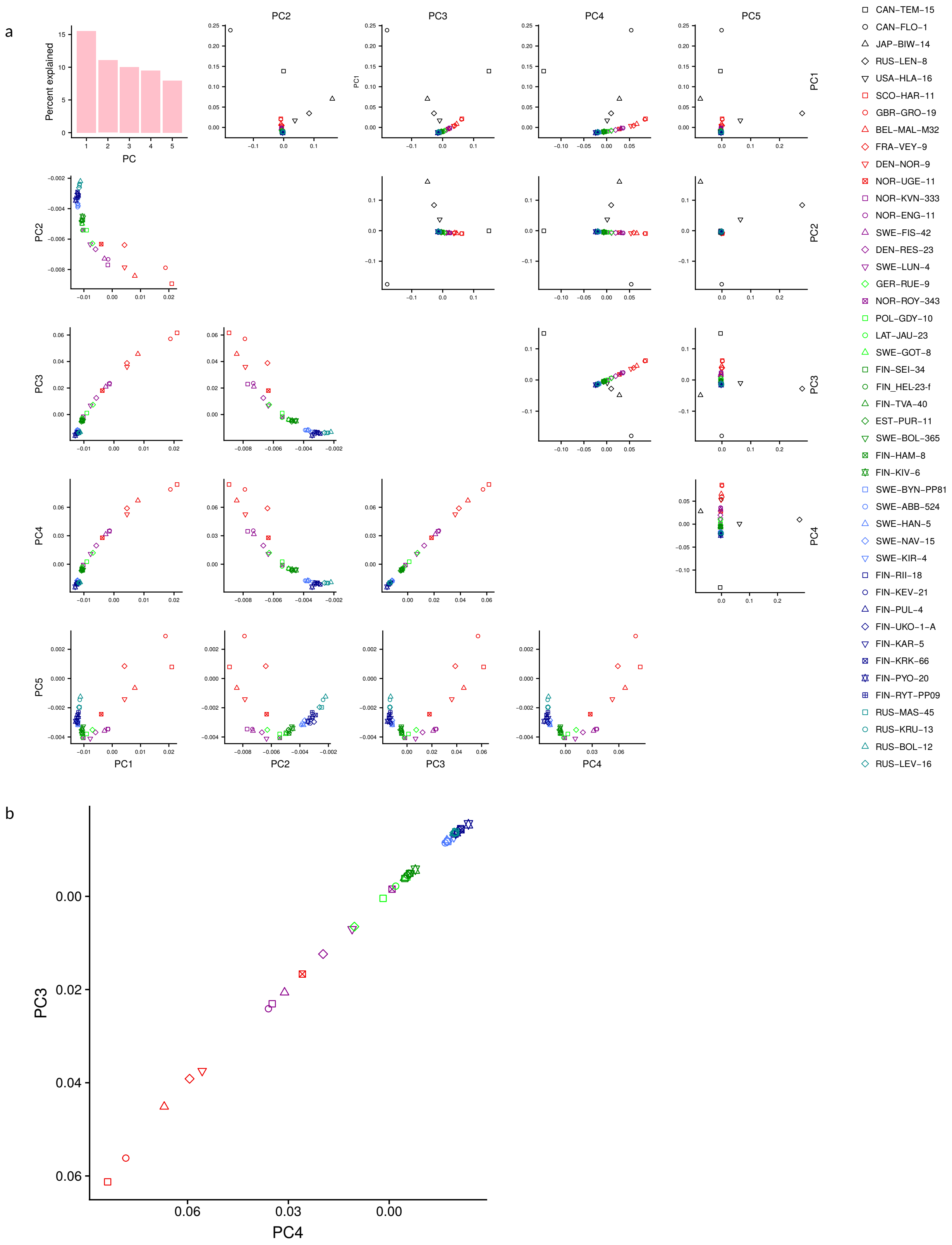


**FIGURE S3** Principal Component Analysis of the full nuclear variant data. a) The subplots show the first five PCA axes for the full data (top) and with the five non-European populations removed (bottom). The barplot shows the proportion of variation explained by each of the PCs. b) After removal of non-European populations, PC3 vs. PC4 shows a continuum from WL to EL with Kattegat/Skagerrak and Baltic populations in between. For clarity, one randomly selected individual from each population is shown.

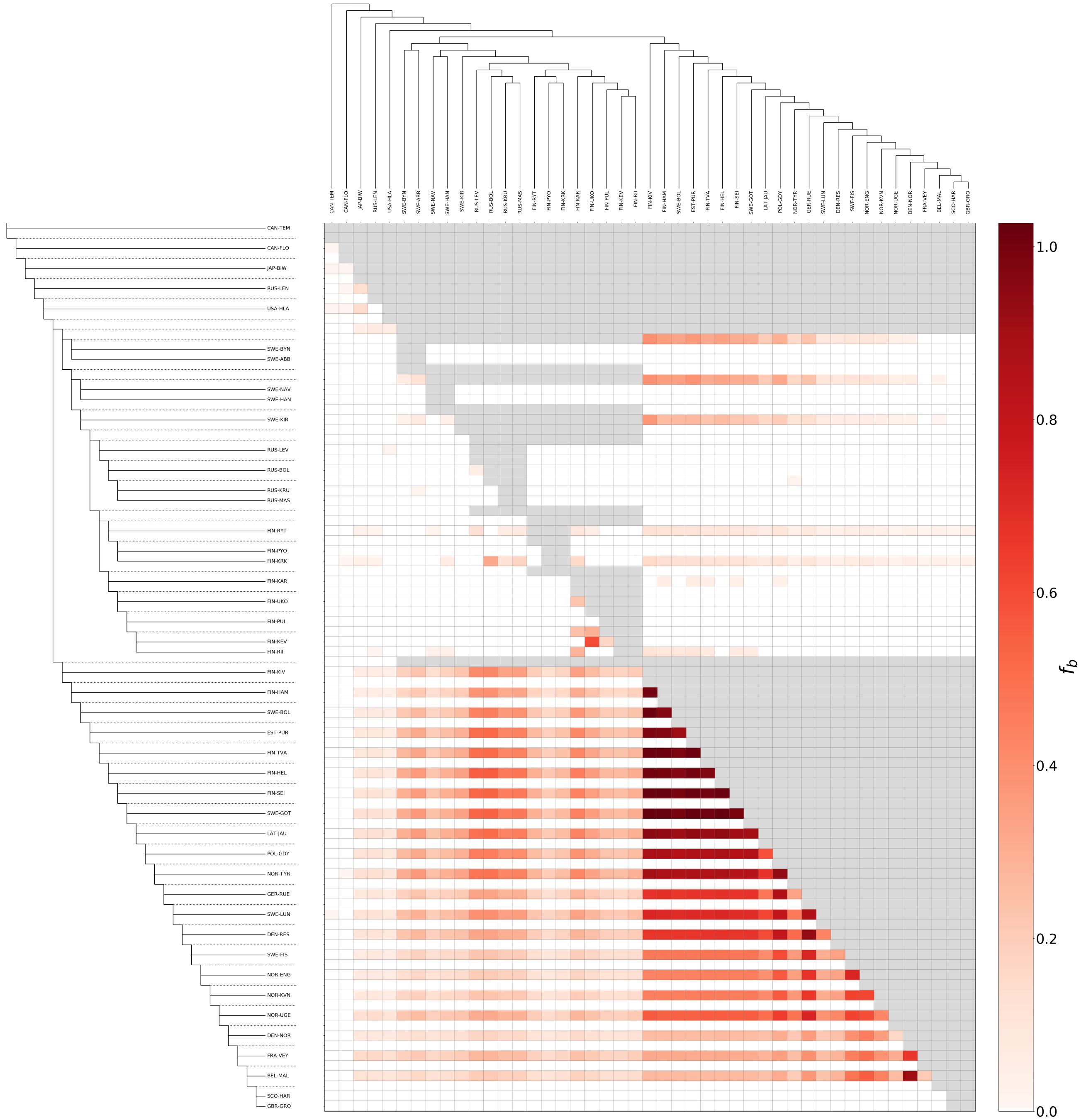


**FIGURE S4** The *f-branch* analysis identifies excess of shared derived alleles between populations (y axis) and the inferred species tree (x axis; ASTRAL tree with *Pungitius tymensis* as the outgroup). The population relationships are based on the frequency of "BBAA" patterns and the matrix values indicate excess of allele sharing between the branches on the y axis and the populations on the x axis, the higher fb values meaning greater allele sharing. All fb values are shown; values marked with gray cannot be estimated based on the given phylogeny.

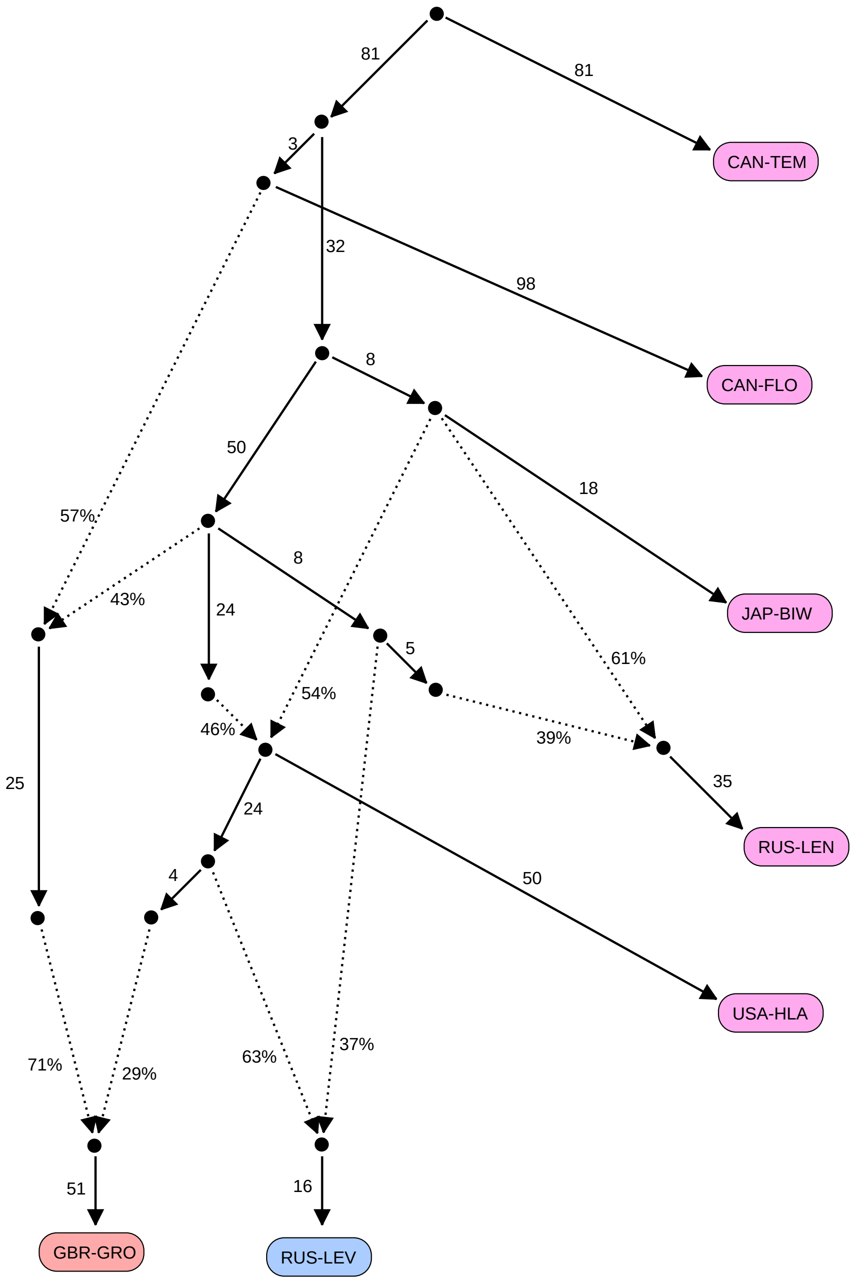


**FIGURE S5** History of North American and Asian populations and representative populations from Europe. The optimal qpGraph model (|Z|= 1.60) places the North American CAN-TEM and CAN-FLO and the Japanese JAP-BIW at the root. The freshwater populations from the River Lena (RUS-LEN) and Alaska (USA-HLA) derive, respectively, 61% and 54% of ancestry from an ancestor of the JAP-BIW, and 39% and 46% from populations representing the later waves of east-to-west migration. The EL population RUS-LEV derives its ancestry from the second and third waves of migration and shares one of these (the graph does not indicate which is earlier) with the WL population GBR-GRO. However, RUS-LEV (and the EL) lacks the old ancestry forming 40% (0.71 x 0.57 = 0.405) of the GBR-GRO genome.

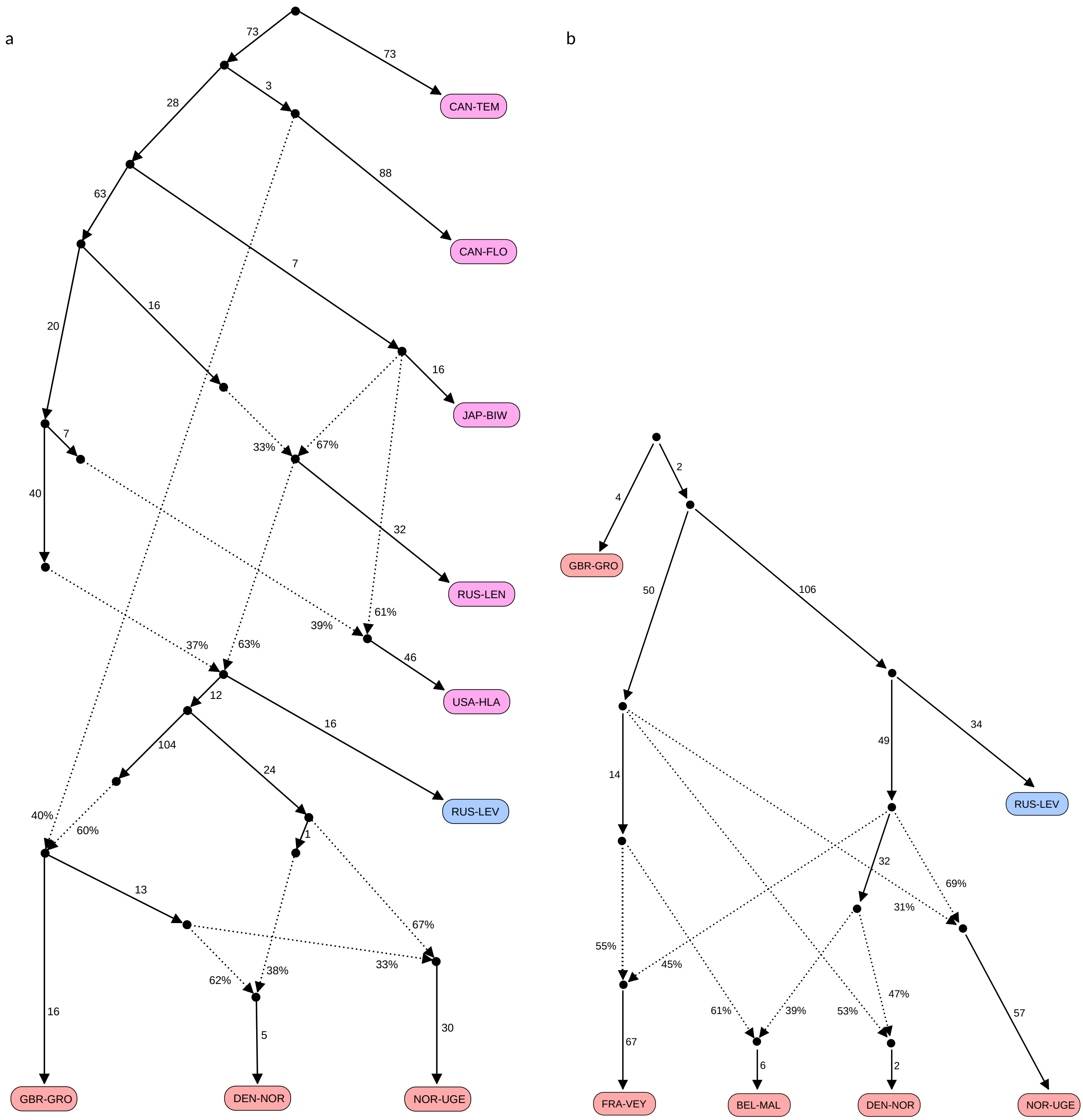


**FIGURE S6** The optimal qpGraph models for non-European populations (magenta) and representatives of WL and EL (red and blue, respectively). a) The full model (|Z|= 2.60) indicates a unique history for the isolated Norwegian population NOR-UGE. The North Sea population from Denmark (DEN-NOR) descends predominantly from the old WL (62%) but has a significant, more recent contribution from the east (38%). The proportions are almost identical to our basal model shown in Fig. 3. NOR-UGE descends predominantly from the east (67%) and has received a lower proportion (33%) from an old WL source shared with DEN-NOR. b) A submodel (|Z|= 2.16) for the European populations indicates a deep divergence of NOR-UGE and a longer shared history for the Belgian (BEL-MAL) and French (FRA-VEY) populations in comparison to DEN-NOR.

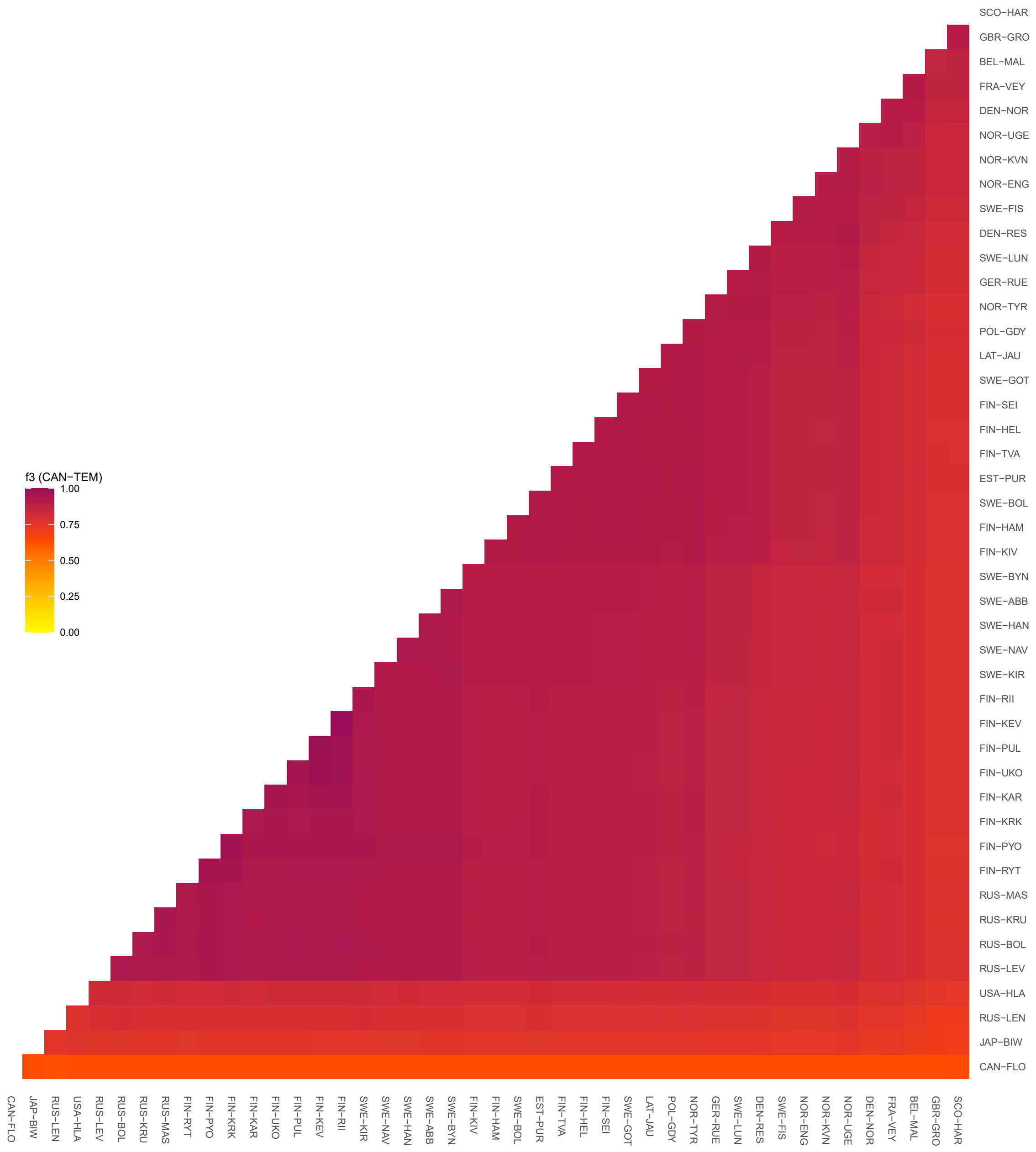


**FIGURE S7** The outgroup-*f3* analysis using CAN-TEM as the outgroup. The heatmap shows the statistics *f3*(X, Y; CAN-TEM) with darker values indicating a closer affinity between populations X and Y.

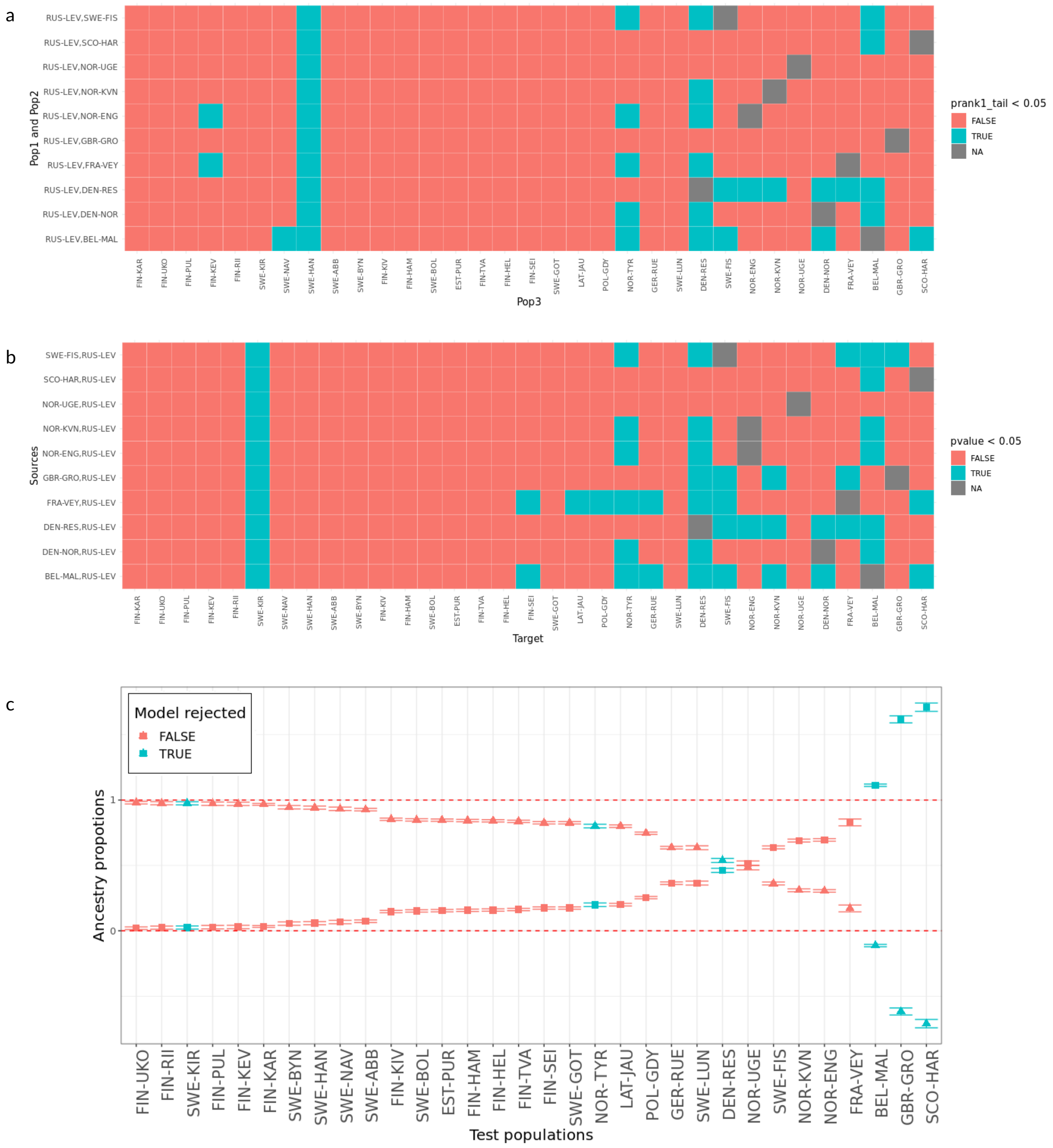


**FIGURE S8** Identification of optimal source pairs for the Baltic Sea populations using qpWave and qpAdm. a) Optimal source pairs for the Baltic Sea populations were identified using qpWave triplet analysis. RUS-LEV was fixed as representative for EL (Pop1). Each WL population (Pop2; y axis) was then tested as the "source" for each of the admixed populations (Pop3; x axis). The ANC5 set was used as the outgroup. The significance of prank1_tail values of each triplet is shown: triplets with insignificant results (p>0.05, red) mean that one of the populations in the triplet can be modeled as a mixture by the other two populations. b) Verification of triplets identified with qpWave using qpAdm. RUS-LEV was fixed as the EL source and the ANC5 set was used as the outgroup. Different WL populations were tested against the target populations. A significant result (p<0.05, shown in cyan) means that the source population pair (y axis) was rejected by the model and is not optimal for modelling the target population (x axis) as two-way admixture. The results indicate that DEN-NOR is the optimal source population for modeling the Baltic Sea populations. c) Admixture proportions with standard errors as estimated by qpAdm. RUS-LEV and DEN-NOR were used as the representatives of the EL and WL sources. Triangles and squares represent the ancestry from EL and WL, respectively. The blue and red color indicates if the model of two-way admixture for the given target population was rejected or not. The red dashed lines refers to admixture proportions of 0 and 1, and estimates beyond the range means the model being rejected. Note that the FRA-VEY population shows a high standard error: although the model was not rejected, it should be considered uncertain (Harney, Patterson, Reich, & Wakeley 2021).

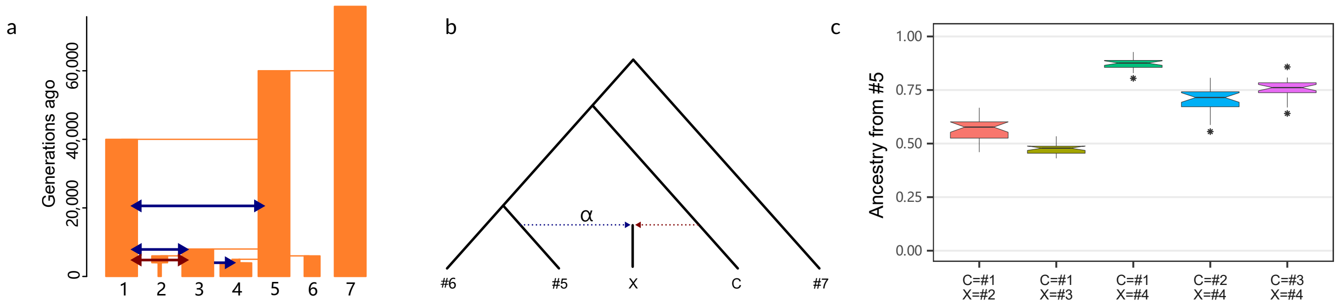


**FIGURE S9** Simulation model and f4-ratio test with synthetic data. a) The simulation model reflects the inferred main events: population #4 represents an admixed Baltic Sea population founded through a bottleneck from an EL population (#5) and subsequently obtaining 25% of its ancestry from a WL population (#3) that itself had a complex history. Largest populations had $N_e$=100,000, other populations are in scale. Blue and red arrows indicate 25% and 15% of the target population(s), respectively, originating from mass migration. b) The setup of the *f4*-ratio test. c) The correct arrangement, C=#3 and X=#4, gives ancestry proportion very close to the true value of 0.75. The use of population #1 and #2 as a source (C=#1, C=#2) for the test population #4 (X=#4) underestimates and overestimates, respectively, the ancestry from population #5. If #1 is used as a source (C=#1), populations #2 and #3 are inferred to have approximately 50% of ancestry from population #5.

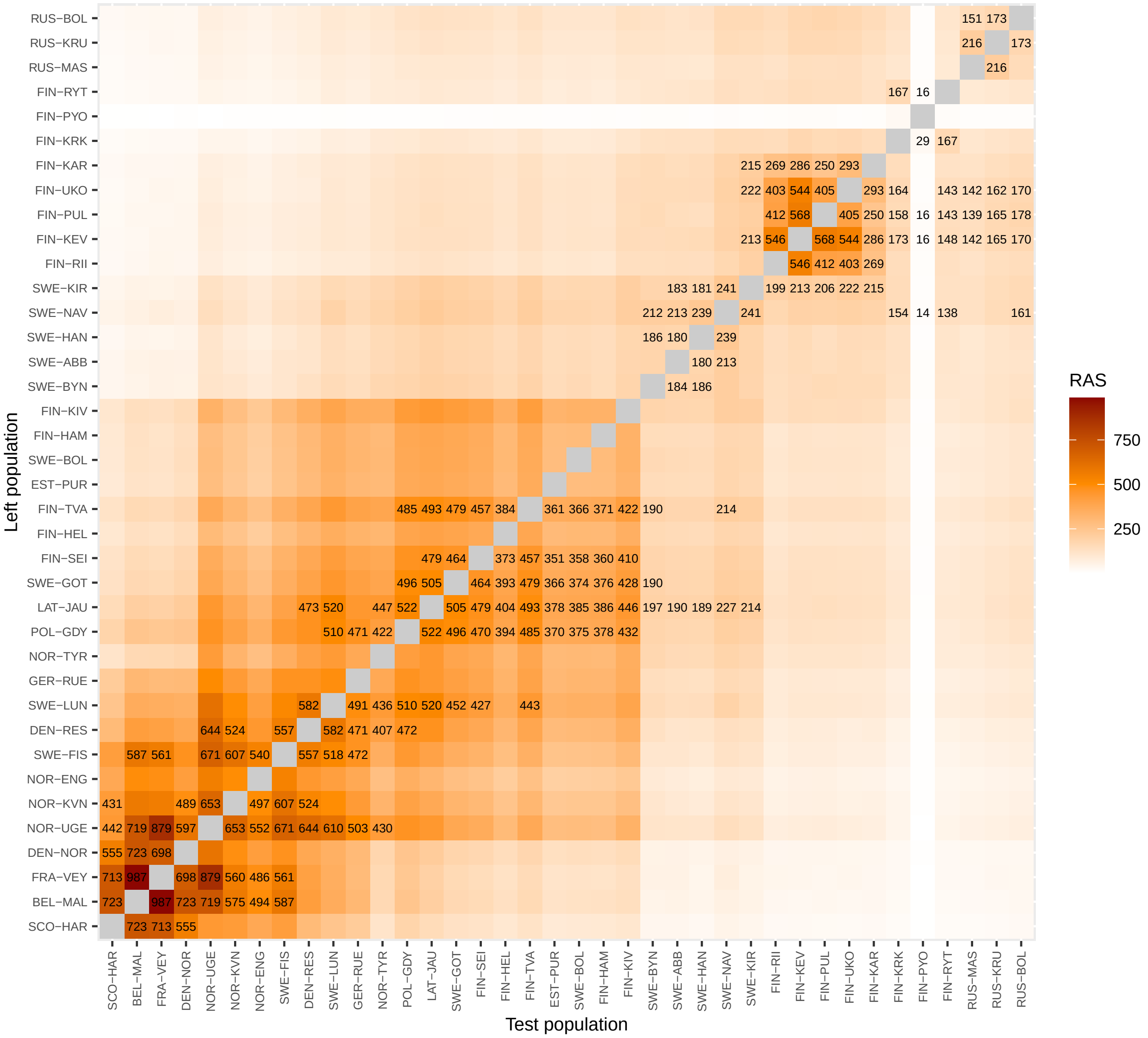


**FIGURE S10** Heatmap of rare alleles sharing statistics (RASS) for each test population. Rare alleles with allele count between 2-10 in the reference set (ANC5, GBR-GRO and RUS-LEV) were included in the statistics. Alleles were polarised using *Pungitius tymensis*. The five highest RASS values for each test population are shown.

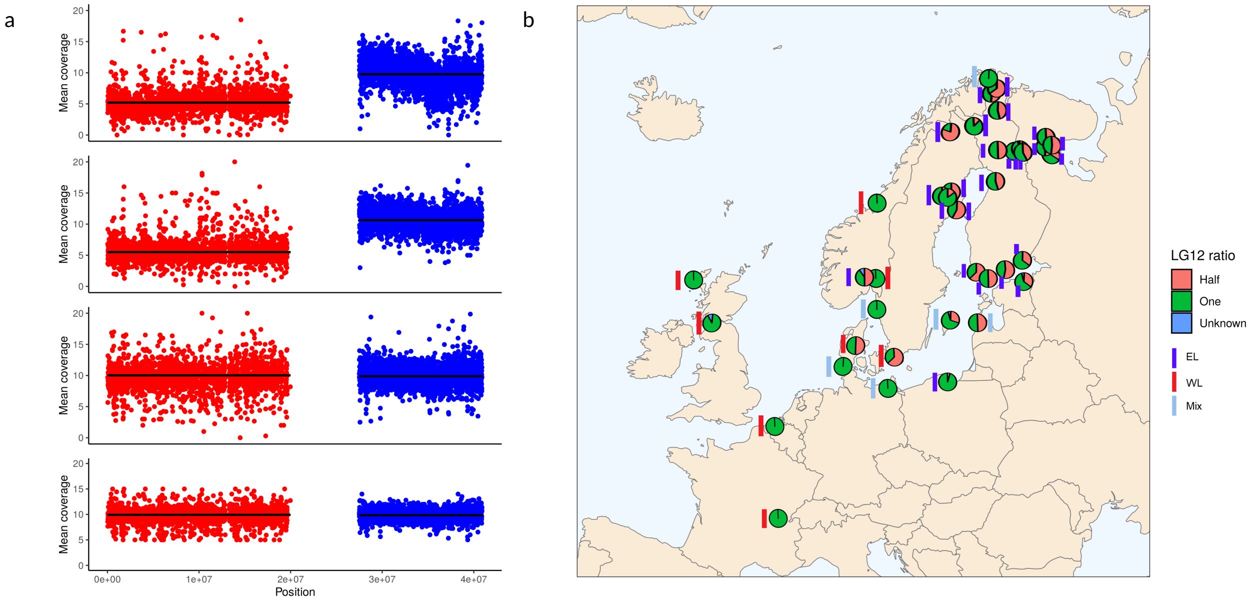


**FIGURE S11** Identification of EL males. a) The mean sequencing coverage across the sex-chromosome (red) and pseudo-autosomal (blue) parts of LG12 in two FIN-PYO males (top) and females (bottom) using 5 kbp windows. The black lines show the average for each region. b) The proportion of samples showing the ratio of half or one for the sequencing coverage across the sex-chromosome and pseudo-autosomal regions. The ratio of half indicates EL male (i.e., EL Y-chromosome) while the ratio of one can either indicate EL female (X-chromosome) or a different sex chromosome system. The colored rectangle indicates the mtDNA type.

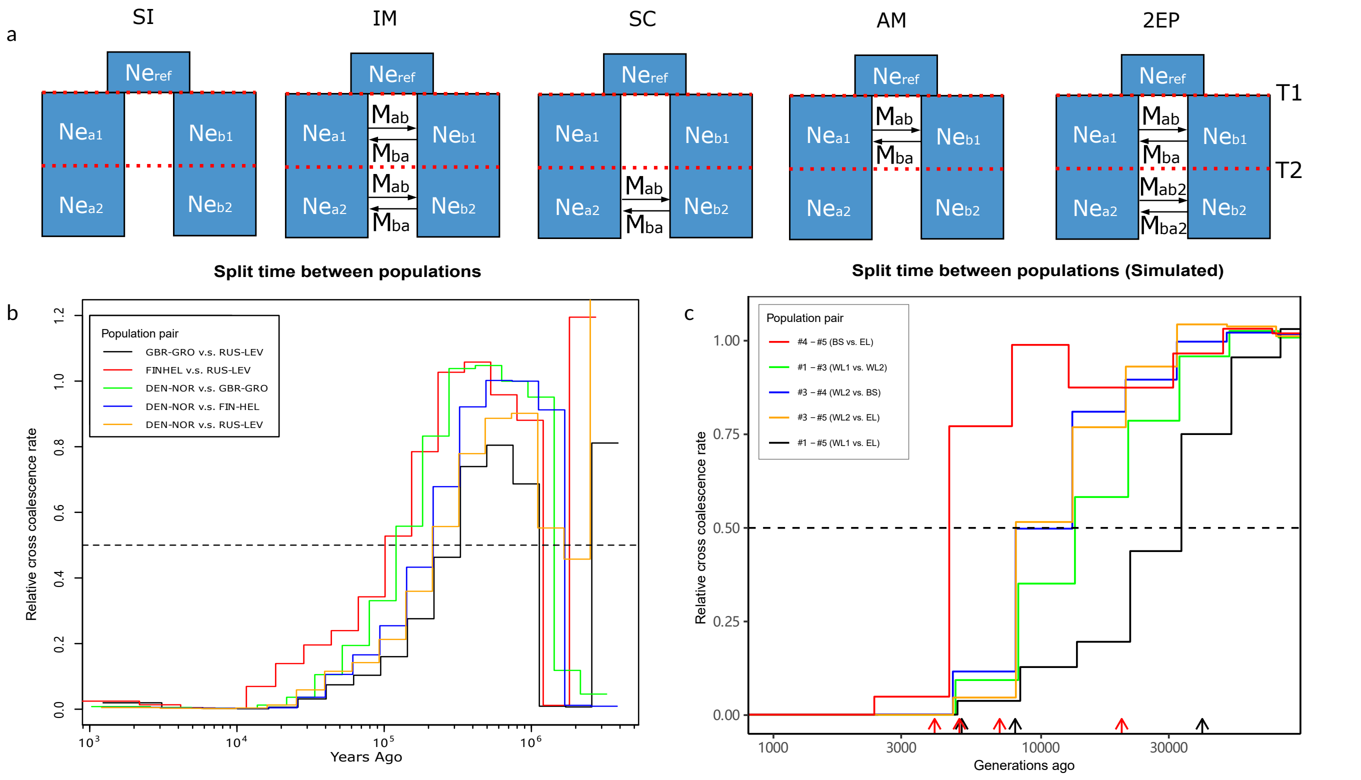


**FIGURE S12** Dating of population separation time with empirical and synthetic data. a) Illustration of the five demographic models used in *moments* analysis: the strict isolation model (SI); the isolation with migration model (IM); the secondary contact model (SC); the ancestral migration model (AM); and the two-epoch model (2EP). The five models allows one change in the $N_e$ at T2. The migration rate were symmetric and fixed in IM, SC and AM model but allowed to change once in the Two-Epoch model at T2. Detailed results of moments analysis for the empirical and synthetic data are shown in Tables S11 and S12, respectively. b) Relative cross-coalescence rates for five population pairs. c) Relative cross-coalescence rates replicated with synthetic data simulated using the model in Fig. 5c. Despite the simplistic scenario, the relative cross-coalescence rate results show the gene flow from ancestral population #1 (“WL 1”) pushing the divergence time e.g. between populations #3 (“WL 2”) and #5 (“EL”) back in time. Black and red arrows at the bottom indicate the times of population divergence and admixture events, respectively. A relative cross-coalescence rate of 0.5 was defined as the divergence time.

**TABLE S1** Sample information, mtDNA type (A-D) and lineage assignment.

| This study (WGS) | | | | | | | | | | | | Teacher et al. 2011 | | Guo et al. 2019 | | |
|---|---|---|---|---|---|---|---|---|---|---|---|---|---|---|---|---|
| | | | | | | mtDNA type | | | | | | | | | | |
| Label | Lat | Long | Habitat | Region | N | A | B | C | D | Assignm. | | Label | Assignm. | Label | Assignm. | Species |
| BEL-MAL* | 51.174 | 3.469 | River | Maldegem; Belgium | 20 | 0 | 0 | 20 | 0 | WL | | BE-LEU | WE | PP-BE-MAL | WE | *P.pungitius* |
| DEN-NOR* | 54.984 | 8.661 | Marine | North Sea; Denmark | 25 | 0 | 11 | 14 | 0 | WL | | | | | | *P.pungitius* |
| DEN-RES | 56.183 | 9.633 | River | Guden; Denmark | 20 | 0 | 0 | 20 | 0 | WL | | DK-GUD | WE | | | *P.pungitius* |
| EST-PUR | 59.416 | 26.983 | River | Purtse River; Estonia | 19 | 18 | 1 | 0 | 0 | EL | | EE-PUR | EE | | | *P.pungitius* |
| FIN-HAM | 60.562 | 27.192 | Marine | Baltic Sea; Finland | 20 | 20 | 0 | 0 | 0 | EL | | | | | | *P.pungitius* |
| FIN-HEL | 60.202 | 25.182 | Marine | Baltic Sea; Finland | 22 | 22 | 0 | 0 | 0 | EL | | FI-HEL | EE | PP-FI-HEL | EE | *P.pungitius* |
| FIN-KAR | 66.656 | 26.440 | Pond | Karhulampi; Finland | 20 | 20 | 0 | 0 | 0 | EL | | FI-KAR | EE | | | *P.pungitius* |
| FIN-KEV | 69.757 | 27.012 | River | Kevojärvi; Finland | 20 | 20 | 0 | 0 | 0 | EL | | FI-KEV | EE | | | *P.pungitius* |
| FIN-KIV | 65.000 | 25.470 | Marine | Baltic Sea; Finland | 19 | 19 | 0 | 0 | 0 | EL | | FI-KIV | EE | | | *P.pungitius* |
| FIN-KRK | 66.437 | 29.135 | Pond | Kirkasvetinenlampi; Finland | 20 | 20 | 0 | 0 | 0 | EL | | FI-KRK | EE | | | *P.pungitius* |
| FIN-PUL | 70.017 | 28.018 | Lake | Pulmankijärvi; Finland | 17 | 17 | 0 | 0 | 0 | EL | | FI-PUL | EE | | | *P.pungitius* |
| FIN-PYO | 66.261 | 29.433 | Pond | Pyöreälampi; Finland | 31 | 31 | 0 | 0 | 0 | EL | | FI-PYO | EE | | | *P.pungitius* |
| FIN-RII | 68.109 | 23.571 | Lake | Riikojärvi; Finland | 23 | 23 | 0 | 0 | 0 | EL | | FI-RIK | EE | | | *P.pungitius* |
| FIN-RYT | 66.384 | 29.320 | Pond | Rytilampi; Finland | 21 | 21 | 0 | 0 | 0 | EL | | FI-RYT | EE | PP-FI-RYT | EE | *P.pungitius* |
| FIN-SEI | 60.233 | 21.954 | Marine | Baltic Sea; Finland | 24 | 24 | 0 | 0 | 0 | EL | | | | | | *P.pungitius* |
| FIN-TVA | 59.833 | 23.200 | Marine | Baltic Sea; Finland | 22 | 22 | 0 | 0 | 0 | EL | | | | | | *P.pungitius* |
| FIN-UKO | 68.776 | 27.436 | Lake | Ukonjärvi; Finland | 22 | 22 | 0 | 0 | 0 | EL | | | | | | *P.pungitius* |
| FRA-VEY* | 46.235 | 5.116 | River | Veyle river; France | 16 | 0 | 0 | 0 | 16 | EL | | | | | | *P.pungitius* |
| GBR-GRO* | 57.615 | -7.511 | River | Loch Grogarry; United Kingdom | 19 | 0 | 0 | 19 | 0 | WL | | | | | | *P.pungitius* |
| GER-RUE* | 54.006 | 13.003 | Marine | Baltic Sea; Germany | 27 | 16 | 1 | 10 | 0 | WL+EL | | GE-RUG | WE+EE | | | *P.pungitius* |
| LAT-JAU | 57.510 | 21.690 | River | Baltic Sea; Lativa | 21 | 19 | 0 | 2 | 0 | EL | | | | | | *P.pungitius* |
| NOR-ENG* | 59.896 | 10.533 | Lake | Engervann; Norway | 23 | 0 | 0 | 23 | 0 | WL | | NO-ENG | WE | | | *P.pungitius* |
| NOR-KVN* | 70.576 | 26.993 | Lake | Kjervatn; Norway | 11 | 0 | 1 | 10 | 0 | WL | | | | | | *P.pungitius* |
| NOR-TYR | 59.910 | 10.299 | Lake | Tyrifjord; Norway | 8 | 8 | 0 | 0 | 0 | EL | | | | | | *P.pungitius* |
| NOR-UGE* | 63.958 | 10.426 | Lake | Ugedalsvatnet; Norway | 20 | 0 | 0 | 0 | 20 | WL | | NO-UGE | WE | PP-NO-UGE | WE | *P.pungitius* |
| POL-GDY | 54.397 | 18.528 | Marine | Baltic Sea; Poland | 20 | 20 | 0 | 0 | 0 | WL+EL | | PL-GDY | EE | | | *P.pungitius* |
| RUS-BOL | 66.295 | 33.366 | Pond | Bolotnoje; Russia | 20 | 20 | 0 | 0 | 0 | EL | | RU-BOL | EE | PS-RU-BOL | EE | *P.pungitius* |
| RUS-KRU | 66.298 | 33.345 | Pond | Krugloje; Russia | 20 | 20 | 0 | 0 | 0 | EL | | RU-KRU | EE | | | *P.pungitius* |
| RUS-LEV | 66.290 | 33.434 | Marine | White Sea; Russia | 30 | 30 | 0 | 0 | 0 | EL | | RU-LEV | EE | PP-RU-LEV | EE | *P.pungitius* |
| RUS-MAS | 66.291 | 33.381 | Pond | Mashinnoje; Russia | 21 | 21 | 0 | 0 | 0 | EL | | RU-MAS | EE | | | *P.pungitius* |
| SCO-HAR* | 55.750 | -4.416 | Lake | Harelaw dam; United Kingdom | 10 | 0 | 0 | 10 | 0 | WL | | UK-HAR | WE | PP-UK-HAR | WE | *P.pungitius* |
| SWE-ABB | 64.478 | 19.436 | Pond | Abborrtjärn; Sweden | 21 | 21 | 0 | 0 | 0 | EL | | SE-ABB | EE | | | *P.pungitius* |
| SWE-BOL | 63.661 | 20.211 | Marine | Baltic Sea; Sweden | 21 | 21 | 0 | 0 | 0 | EL | | SE-BOL | EE | | | *P.pungitius* |
| SWE-BYN | 64.455 | 19.444 | Pond | Bynästjärnen; Sweden | 23 | 23 | 0 | 0 | 0 | EL | | SE-BYN | EE | PP-SE-BYN | EE | *P.pungitius* |
| SWE-FIS* | 58.233 | 11.400 | Marine | Fiskebackskil; Sweden | 20 | 1 | 0 | 19 | 0 | WL+EL | | SE-FIS | WE+EE | PP-SE-FIS | WE | *P.pungitius* |
| SWE-GOT | 57.733 | 18.950 | Marine | Baltic Sea; Sweden | 19 | 18 | 0 | 1 | 0 | WL+EL | | | | | | *P.pungitius* |
| SWE-HAN | 64.556 | 19.173 | Pond | Hansmyrtjärn; Sweden | 20 | 20 | 0 | 0 | 0 | EL | | SE-HAN | EE | | | *P.pungitius* |
| SWE-KIR | 67.896 | 20.086 | Pond | Kiruna; Sweden | 15 | 15 | 0 | 0 | 0 | EL | | | | | | *P.pungitius* |
| SWE-LUN | 55.716 | 13.433 | Pond | Baltic Sea; Sweden | 21 | 0 | 0 | 21 | 0 | WL+EL | | SE-LUN | WE | | | *P.pungitius* |
| SWE-NAV | 64.564 | 19.198 | Pond | Lil-Navartjärn; Sweden | 20 | 20 | 0 | 0 | 0 | EL | | SE-NAV | EE | | | *P.pungitius* |
| CAN-FLO | 54.233 | -111.633 | Lake | Floatingstone Lake; Canada | 8 | | | | | ANC | | | | PP-CA-FLO | NA | *P.pungitius* |
| CAN-TEM | 47.713 | -68.916 | Lake | Lac Témiscouata; Canada | 16 | | | | | ANC | | CA-TAM | CA(NA) | PP-CA-TEM | NA | *P.pungitius* |
| JAP-BIW | 43.082 | 145.113 | Marine | Biwase Bay; Japan | 24 | | | | | ANC | | | | PP-JP-BIW(BW) | FE | *P.pungitius* |
| RUS-LEN | 72.983 | 126.966 | River | Lena River; Russia | 10 | | | | | ANC | | | | PP-RU-LEN | FE | *P.pungitius* |
| USA-HLA | 61.590 | -149.760 | Lake | Hawks Lane; United States | 20 | | | | | ANC | | | | | | *P.pungitius* |
| PUN-TYM | 43.827 | 145.086 | River | Motosakimui River; Japan | 1 | | | | | Outgroup | | | | PT-JP-MOT | Outgroup | *P.tymensis* |

* Populations where the EL male (XY) sex-chromosomes not presenting.

**TABLE S2** Data filtering for different analyses.

| Name | Filtering criteria | SNPs | Analyses used |
|---|---|---|---|
| SNP Set 0 | Of original dataset, keep autosomal biallelic SNPs in non-repetitive regions; remove interspecific variants | | *NA* |
| SNP Set 1 | From SNP Set 0, keep only binary SNPs with quality score $\geq$30, mean coverage $\geq$8x, GQ $\geq$ 20 | 152,175,77 | *NA* |
| SNP Set 2 | From SNP Set 1, filter in VCFtools with --minGQ 20 --minQ 30 --min-meanDP 8 --max-meanDP 25 --max-missing 0.9, and then LD pruning with plink --indep-pairwise 50 10 0.1 | | PCA, ADMIXTURE |
| SNP Set 3 | From SNP Set 1, select two random samples from each populations allowing maximum 50% missing data and keep only variable sites with BCFtools view (--min-ac=1) | | RAxML and ASTRAL |
| SNP Set 4 | From SNP Set 1, keep only variable sites with maximum mean DP of 25 and no more than 25% missing data; distance between SNPs required to be at least 300bp | 506,171 | ADMIXTOOLS (f4 statistic, outgroup-*f3*, *f4*-ratio, qpWave, qpAdm and qpGraph test) and RASS |
| SNP Set 5 | From SNP Set 1, keep only variable sites with maximum mean DP of 25 and no more than 10% missing data | | *f-branch* |
| | For details, see https://github.com/XueyunF/nsp_phylogeo/blob/main/Divergence_time/MSMC2_pipeline.md | | MSMC2 |
| | For details, see https://github.com/XueyunF/nsp_phylogeo/blob/main/Divergence_time/Moments_pipeline.md | | *moments* |
| SNP Set 6 | Subsets of the EL sex chromosome (LG12). For details, see https://github.com/XueyunF/nsp_phylogeo/blob/main/Sex_chromosome/README.md | | EL sex determination |
| SNP Set 7 | Complete mitochondrial genome. For details, see https://github.com/XueyunF/nsp_phylogeo/tree/main/Structure_and_Phylogeny | | RAxML |

**TABLE S3** Detailed results of qpWave analysis: WL, EL and Baltic Sea populations.

| | Left (target) populations | Right (source) populations | p_rank0 | p_rank1 | p_rank2 | p_rank3 | p_rank4 | p_rank5 | p_rank6 |
|---|---|---|---|---|---|---|---|---|---|
| Test WL | GBR-GRO, SCO-HAR, DEN-NOR, FRA-VEY, NOR-UGE, BEL-MAL, NOR-KVN, SWE-FIS, NOR-ENG | CAN-TEM, CAN-FLO, JAP-BIW, RUS-LEN, USA-HLA, RUS-LEV | 0 | 2.86E-60 | 0.209 | 0.681 | 0.89 | 1 | NA |
| Test EL | RUS-LEV, RUS-KRU, RUS-MAS, RUS-BOL, FIN-KRK, FIN-PYO, FIN-RYT, FIN-UKO, FIN-RII, FIN-PUL, FIN-KEV, FIN-KAR, SWE-ABB, SWE-HAN, SWE-BYN, SWE-NAV, SWE-KIR | CAN-TEM, CAN-FLO, JAP-BIW, RUS-LEN, USA-HLA, GBR-GRO | 7.98E-94 | 3.33E-13 | 0.003 | 0.829 | 0.998 | 1 | NA |
| Test BS | FIN-KIV, FIN-HAM, SWE-BOL, EST-PUR, FIN-TVA, FIN-HEL, FIN-SEI, SWE-GOT, LAT-JAU, POL-GDY, GER-RUE, SWE-LUN | CAN-TEM, CAN-FLO, JAP-BIW, RUS-LEN, USA-HLA, GBR-GRO, RUS-LEV | 0 | 2.56E-66 | 0.071 | 0.891 | 0.966 | 0.835 | 1 |
| Test BS* | FIN-KIV, FIN-HAM, SWE-BOL, EST-PUR, FIN-TVA, FIN-HEL, FIN-SEI | CAN-TEM, CAN-FLO, JAP-BIW, RUS-LEN, USA-HLA, GBR-GRO, RUS-LEV | 4.4E-162 | 0.144 | 0.891 | 0.878 | 0.993 | 0.791 | 1 |

*Excluding southern Baltic Sea populations and LAT-JAU

**TABLE S4** Detailed results of qpWave analysis: WL and EL populations.

| | Left (target) populations | | Results | | | Right (source) populations |
|---|---|---|---|---|---|---|
| | FIX | TEST | p_rank0 | p_rank1 | p_rank2 | |
| Within WL | GBR-GRO, SCO-HAR | DEN-NOR | <0.001 | 0.024 | 1 | ANC5, RUS-LEV |
| | | BEL-MAL | <0.001 | 0.07 | 1 | |
| | | FRA-VEY | <0.001 | 0.089 | 1 | |
| | | NOR-UGE | 0 | 0.022 | 1 | |
| | | NOR-KVN | <0.001 | 0.001 | 1 | |
| | | SWE-FIS | 0 | 0.004 | 1 | |
| | | NOR-ENG | 0 | 0.009 | 1 | |
| | | DEN-RES | 0 | 0.004 | 1 | |
| Within EL | RUS-LEV, RUS-BOL | RUS-KRU | 0.151 | 0.152 | 1 | ANC5, GBR-GRO |
| | | RUS-MAS | 0.595 | 0.628 | 1 | |
| | | FIN-RYT | 0.891 | 0.984 | 1 | |
| | | FIN-PYO | 0.802 | 0.867 | 1 | |
| | | FIN-KRK | 0.507 | 0.356 | 1 | |
| | | FIN-UKO | 0.182 | 0.411 | 1 | |
| | | FIN-PUL | 0.070 | 0.413 | 1 | |
| | | FIN-KEV | 0.025 | 0.418 | 1 | |
| | | FIN-KAR | <0.001 | 0.407 | 1 | |
| | | SWE-ABB | <0.001 | 0.442 | 1 | |
| | | SWE-HAN | <0.001 | 0.418 | 1 | |
| | | SWE-BYN | <0.001 | 0.392 | 1 | |
| | | SWE-NAV | <0.001 | 0.441 | 1 | |
| | | SWE-KIR | 0.020 | 0.373 | 1 | |

**TABLE S5** Detailed results of qpWave analysis: WL populations.

| Left (target) populations | | Results | | | | | Right (source) populations |
|---|---|---|---|---|---|---|---|
| FIX | TEST | p_rank0 | p_rank1 | p_rank2 | p_rank3 | p_rank4 | |
| GBR-GRO, SCO-HAR, BEL-MAL, FRA-VEY | - | <0.001 | 0.228 | 0.713 | 1 | NA | ANC5, RUS-LEV |
| | DEN-NOR | <0.001 | <0.001 | 0.948 | 0.9 | 1 | |
| | NOR-UGE | 0 | <0.001 | 0.820 | 0.785 | 1 | |
| | NOR-KVN | 0 | <0.001 | 0.843 | 0.965 | 1 | |
| | SWE-FIS | 0 | <0.001 | 0.877 | 0.891 | 1 | |
| | NOR-ENG | 0 | <0.001 | 0.846 | 0.969 | 1 | |
| | DEN-RES | 0 | <0.001 | 0.844 | 0.892 | 1 | |
| GBR-GRO, SCO-HAR, BEL-MAL, FRA-VEY | - | <0.001 | 0.744 | 0.973 | 1 | NA | ANC5 |
| | DEN-NOR | <0.001 | 0.006 | 0.972 | 0.699 | 1 | |
| | NOR-UGE | <0.001 | 0.053 | 0.801 | 0.811 | 1 | |
| | NOR-KVN | <0.001 | 0.026 | 0.941 | 0.898 | 1 | |
| | SWE-FIS | <0.001 | <0.001 | 0.976 | 0.685 | 1 | |
| | NOR-ENG | <0.001 | 0.007 | 0.945 | 0.857 | 1 | |
| | DEN-RES | <0.001 | <0.001 | 0.839 | 0.862 | 1 | |

**TABLE S6** Two population symmetrical test with qpWave, the source was set as ANC5.

Table S6 is available at https://github.com/XueyunF/nsp_phylogeo/blob/main/Tables/Two_pop_symmetrical_qpwave.txt

**TABLE S7** Detailed results qpWave testing of triples for the Baltic sea populations.

Table S7 is available at https://github.com/XueyunF/nsp_phylogeo/blob/main/Tables/Baltic_triplets_qpWave.txt

**TABLE S8** Detailed results of *f4*-ratio test with the EL as source.

Table S8 is available at https://github.com/XueyunF/nsp_phylogeo/blob/main/Tables/f4_ratio_sourceEL.txt

**TABLE S9** Detailed results of *f4*-ratio test with the WL as source.

Table S9 is available at https://github.com/XueyunF/nsp_phylogeo/blob/main/Tables/f4_ratio_sourceWL.txt

**TABLE S10** Detailed results of f4-statistics.

Table S10 is available at https://github.com/XueyunF/nsp_phylogeo/blob/main/Tables/Dstat_result_full_table.txt.

**TABLE S11** Detailed results of moments analysis for the empirical data.

Table S11 is available at https://github.com/XueyunF/nsp_phylogeo/blob/main/Tables/moments_Emprical.csv.

**TABLE S12** Detailed results of moments analysis for the synthetic data.

Table S12 is available at https://github.com/XueyunF/nsp_phylogeo/blob/main/Tables/moments_Synthetic.csv.

**SCRIPT S1** Python-code for demographic analyses with *moments*

The code is available at https://github.com/XueyunF/nsp_phylogeo/tree/main/Divergence_time.

**SCRIPT S2** Python-code for data simulation with *msprime*

The code is available at https://github.com/XueyunF/nsp_phylogeo/tree/main/Simulations.
